# GaussF: An Unbiased Gaussian Framework for High-Precision Transcript Quantification with a Programmable Self-Benchmark Model

**DOI:** 10.64898/2026.10.28.762697

**Authors:** Qiang Su, Yi Long, Deming Gou, Junmin Quan, Qizhou Lian

## Abstract

Short-read RNA sequencing has revolutionized transcriptome analysis, but accurate isoform-level quantification faces substantial challenges from overlapping sequences among complex isoforms and inherent sequencing biases, leading to uneven coverage. Current alignment-based and alignment-free methods, while valuable, largely rely on empirical, data-derived factors for bias correction and isoform resolution. This dependency limits their capacity to fully capture the intricate spectrum of sequencing data. We introduce GaussF, a novel alignment-free methodology specifically designed for precise isoform quantification by directly addressing these biases and the isoform recognition problem without relying on empirical data. GaussF is predicated on the observation that k-mer counts categorized by GC content across a natural transcript exhibit an intrinsic Gaussian distribution. It establishes a transcript-specific self-benchmarking workflow utilizing a theoretical Gaussian cumulative distribution function (CDF). Crucially, the parameters defining this CDF are derived independently of any empirical sequencing data. This parameter-fixed theoretical CDF serves as an unbiased benchmark, capturing only the genuine contribution of abundance while systematically excluding all bias-related data. To improve isoform resolution despite shared regions, GaussF characterizes isoforms through identifying minimal-shared regions and restructures the k-mer reference accordingly. This approach simplifies complexity and enhances accuracy. Rigorous validation using artificial and real RNA samples, alongside comparative analysis with leading quantification tools like Kallisto, Salmon, Cufflinks, StringTie, and RSEM, demonstrates GaussF’s superior performance in accurate isoform detection and quantification. GaussF is available on GitHub and PyPI, representing a significant advancement in RNA-seq analysis.

## Introduction

RNA sequencing (RNA-seq) has revolutionized our comprehension of gene expression and regulation, granting unprecedented insight into the dynamic realm of transcriptomes.^1, 2^ This powerful technique has become an indispensable tool for dissecting diverse biological phenomena by accurately quantifying transcript abundances, especially at isoform level, from unraveling intricate cellular processes to elucidating disease mechanisms.^3, 4^ Currently, RNA-seq largely employs short-read sequencing for its data acquisition phase, chosen for its high-throughput capabilities and exceptional data quality.^5–7^ This method excels due to parallelization and sophisticated error correction, which together allow for a detailed representation of the transcriptome encompassing even the scarcest transcripts. High-caliber data is essential for the intricate analysis that follows, including precise transcript quantification. Nevertheless, the full value of RNA-seq, especially when leveraging short-read technology, hinges on a rigorous empirical workflow.^8, 9^ This combination ensures the successful translation of undeciphered sequence information from long, intact transcript molecules into multiple short, readable fragments. Despite these notable effectiveness, the accurate quantification of transcripts, in particular at isoform level, from numerous short fragments poses substantial challenges. The complete sequence of any given isoform typically overlaps with sequences from multiple other complex isoforms. This overlapping nature results in varying levels of sequencing coverage across different segments. Furthermore, inherent biases in sequencing add another layer of complexity, causing coverage to be uneven and highly variable. As a result, all transcript isoforms that undergo sequencing face these dual challenges: the uneven coverage due to the overlap with other isoforms, and the inconsistent coverage stemming from intrinsic sequencing biases.^10^

The pursuit of accurate transcript quantification depends on the employment of either alignment-based or alignment-free methodologies with common maximization algorithm to gauge individual RNA transcript abundance. Alignment-based methods involve the alignment of short RNA-seq reads to a reference genome or transcriptome with the aid of sophisticated algorithms such as Bowtie2,^11^ STAR,^12^ or HISAT2.^13^ Following alignment, read counts or expression values are derived by attributing each read to distinct genomic features. This is accomplished at either gene-level utilizing tools like featureCounts,^14^ or at the isoform-level employing software such as Cufflinks,^15^ StringTie,^16^ and RSEM.^17^ With its well-established practices and long-standing use in the field, these approaches offer a quantitative outlook on the transcriptome, thereby enhancing our comprehension of genomic functions. However, alignment-based approaches may encounter challenges in mapping uncertainty due to alternative splicing events and have limitations from strong memory-required setups.^18^ Conversely, alignment-free methodologies bypass the necessity for read alignment to a reference, opting instead for capturing sequence compositions of the reads. For example, k-mer based algorithms such as Kallisto^19^ and Sailfish^20^ dissect the sequences into fixed-length subsequences (k-mers) and catalog the occurrences of each k-mer throughout the respective dataset. Other alignment-free algorithms, like Salmon,^21^ utilize graph-based representations to extract k-mer features from the reads. These methods offer streamlined, adaptable, and light-footprint solutions to transcript quantification in diverse research scenarios. However, the precision of alignment-free transcriptome assembly quantification remains a challenge due to factors such as sequence complexity, algorithm computational limitations, and isoform resolution.^22^ While it was traditionally believed that alignment-based genome-guided assembly methods offered superior accuracy,^16^ recent studies indicate that alignment-free methods can achieve comparable or even superior results, especially in full-length isoform quantification.^23^ The degree of precision attained in transcript quantification, through both alignment-based and alignment-free methodologies, is fundamentally intertwined with the effectiveness of strategies employed for specific isoform identification and the mitigation of various biases. Central to all current methodologies is the use of predefined factors, which are empirically derived from sequencing data, applied proactively. Critical to understand is that these factors, including isoform likelihood and bias correction coefficients, are determined and utilized in a tailored manner, aimed at addressing specific scenarios with precision. However, this dependency on factors derived from empirical data poses a considerable challenge, limiting their capacity to fully parse the complex spectrum of sequencing data.

In our investigation, we introduce GaussF, a cutting-edge methodology designed to accurately recognize the isoforms and address all sources of biases within an alignment-free framework, thereby facilitating precise isoform-level transcript quantification from the short read sequencing data. Our strategic exploit stems from the observation that the quasi-random distribution of guanine (G) and cytosine (C) bases across a natural transcript engenders an intrinsic Gaussian distribution of k-mer counts that are categorized by the GC content of k-mers. Accordingly, we construct a pseudo-cumulative distribution function (pseudo-CDF) by interpreting the pseudo-Gaussian distribution. This distribution function facilitates a transcript-specific self-benchmarking workflow, underpinned by a dual-distribution model: a theoretically derived uniform distribution of modeling k-mers and an empirically observed, uneven distribution stemming from sequencing kmers across the transcript length. Central to this methodology are the concepts of categorizing k-mers, aggregating GC-indexed k-mer counts, and fitting GC-categorized k-mer counts distribution. The initial phase establishes a firm groundwork for a transcript-specific evaluation of modeling data characterized by a uniform k-mer distribution. It involves an investigation of aggregated k-mer counts (frequencies) of GC-specific k-mers. Subsequently, the count (frequency) aggregates for distinct GC-content categories are estimated through a Gaussian distribution function. In parallel, the cumulated count (frequency) aggregates across these categories are modeled utilizing a CDF of a Gaussian model. The effectiveness of this analytical approach hinges on the precise definition of essential parameters, especially the mean and standard deviation indicative of each transcript’s unique distribution characteristics. The sequencing data are also categorized and aggregated based on their GC content. The GaussF model fits the cumulative sequencing k-mer count aggregates across GC categories using a Gaussian CDF, determined by these predefined parameters. The resulting amplitude form CDF fitting serves as an unbiased abundance indicator, accurately mirroring the total sequencing-count contributions of individual k-mers across the transcript. Characterized as a self-benchmarking process, GaussF’s methodology surpasses the limitations of traditional empirical and/or statistical methods by reaching a theoretical benchmark for joint bias adjustment, rather than merely approximating individual bias. Significantly, the core pre-determined parameters are entirely independent of empirical sequencing data. Consequently, the parameter-fixed Gaussian CDF exclusively captures the genuine contribution of abundance while systematically excluding all bias-related data. This methodology thereby guarantees a unbiased quantification. However, the accurate attribution of sequencing reads to specific isoforms is obstructed by regions shared across isoform variants for sequencing count acquisition. Due to the fact that various segments within an isoform’s complete sequence are shared across a multitude of complex isoforms, the uneven coverage resulting from inherent bias complicates the precise assignment of sequencing reads or k-mer counts to distinct isoforms. To effectively navigate this challenge, certain methodologies have been developed that leverage the utilization of regions characterized by definitive overlap for the precise demarcation of specific isoforms.^24–26^ Inspired by these advances, the GaussF method is predicated on the characterization of isoforms through regions minimally shared among them, employing two principal strategies: the identification of unique regions for single isoform and the determination of minimal-shared regions for multiple overlapping isoforms. The former pertains to isolating a single isoform that exhibits unique regions unshared with others, serving as a definitive representation for that specific isoform. The latter involves identifying a set of more than one isoform that collectively present the least amount of shared regions, thereby representing such overlapping isoforms with minimal ambiguity. The complexity of overlapping isoforms could lead to the scattering of minimal-shared regions across the entire transcript. To tackle these complexities, GaussF approach programmablly restructures the k-mer reference database by integrating these separated minimal-shared regions, thus setting a more robust benchmark for quantification. The methodology presented here simplifies complexity and lowers computational demands efficiently. GaussF is easily accessible through our GitHub repository and is also available on PyPI for straightforward installation. To assert GaussF’s validity, we implemented a rigorous testing protocol that merges theoretical frameworks with empirical tests. This in-depth validation includes experiments using both artificial RNA constructs and real human tissue RNA samples. We also conducted comparative tests against leading transcript quantification methodologies, such as alignment-based and alignment-free approaches like Kallisto, Salmon, Cufflinks, StringTie, and RSEM, to demonstrate GaussF’s enhanced performance in isoform detection and quantification. This extensive validation process affirms GaussF’s significant improvements in RNA-seq data analysis.

## Results

### Theoretical framework and empirical validation of the GC content-dependent GaussF algorithm

The binomial distribution is a statistical model characterizing the probability of achieving a distinct number of successes in a set number of independent trials. It is notably valuable in the appraisal of GC content within a given bin size, k, of RNA fragment. Within this framework, the trial in each position along RNA fragment manifests only two potential outcomes: success (embodied by a degenerate base S-either G or C), or failure (represented by a base W-A or U). Here, the likelihood of encountering a degenerate base S in each bin position is denoted by p, while the possibility of observing a base W equates to q = 1 - p. Since the content of G and C in a sequence could range from 0 to the sequence length, k, the types of GC content possible in the sequences would be k+1. This is because the proportion of G and C could vary anywhere from all nucleotide positions being A or U (giving 0 G or C, hence a GC content type of 0), to all positions being G or C (giving a GC content type of k). So, the total number of GC content types is k + 1. To deduce the binomial distribution for GC content, a simplified model is drawn upon for RNA fragments at a fixed length. This model considers a sequence of n independent Bernoulli trials across the RNA fragment, where each trial ascribes a GC base with a success probability of p = 1/2. Let X denote the random variable reflecting the count of successful GC ascriptions within these n trials (n=k). To ascertain the probability mass function (PMF) of X, the likelihood of attaining X successes is determined. For each success, the success probability (p) is multiplied by itself X times (p^X^). Similarly, for each failure, the failure probability (1-p) is multiplied by itself (n-X) times ((1-p)^(n-X)^). The total combinations of arranging X successful GC-ascriptions amongst the n trials are represented by the binomial coefficient, signifying the possible selection methods of X items from n.

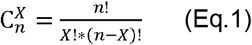

The PMF of X is the product of these three terms:

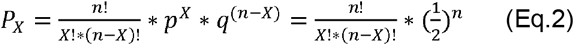

Utilizing the binomial distribution allows for an accurate representation of GC-content distribution characteristics of RNA fragment at a fixed length, attainable through the quasi-random permutation of G and C bases. Nonetheless, it is crucial to acknowledge that the binomial distribution, with its predefined parameters, is optimally suited for illustrating outcomes of unique but identical trials with an unchanging number of trials (n), denoting the fixed RNA length. However, it falls short in scenarios involving variable RNA mixtures or other sources of unpredictability. To circumvent this restriction, we have extracted a flexible Gaussian function from the binomial distribution function. When the values of n (the number of trials), np (the expected number of successes), and nq (the expected number of failures) are sufficiently large, the binomial distribution can be closely approximated by the Gaussian distribution, according to de Moivre’s Laplace limit theorem.^27, 28^ This function is detailed as follows:

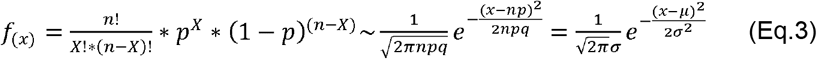

The Gaussian distribution function provides an elevated level of flexibility and practicality in data analysis, offering accommodation for a wide array of scenarios beyond the restrictions imposed by the binomial distribution. As a result, it enables a more precise and versatile analysis of GC distribution patterns across an assortment of RNA fragments.

Building on this concept, we have adjusted the Gaussian distribution function to represent frequency or occurrence, instead of just probability. This adaptation translates the PMF into an occurrence mass function (OMF). To implement this, the PMF is multiplied by the total number of observations ‘A’ within the transcript-specified data set. In this context, the total number of observations is understood to be the sum of categorized GC k-mers. This modified function then reads as follows:

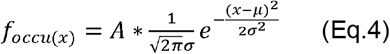

In refining the function to accurately mirror actual data occurrences, the practical utility of our calculations is significantly amplified. This has been facilitated through the development of a cumulative function, which is based on the modified OMF using mathematical integration, with a distinct focus on GC content. Here, ‘f_occu(x)_’ is representative of the occurrence of distinct GC content ‘x’. In view of this, the complete gamut of GC content from the lowest (GC_min_) to the highest (GC_max_) is considered, with the accumulation of these occurrences derived through integration. This mathematical process is encapsulated in the equation as follows:

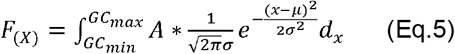

The cumulative function is denoted by F(X) and corresponds to the integral sum of all individual occurrence for every value of GC content ‘x’, thereby yielding a cumulative function that provides the total occurrence of GC contents up to a specific value ‘X’.

To establish a self-benchmarking model for transcript quantification, termed GaussF, we leverage GC-content categories. We first constructed a synthetic RNA-seq dataset based on random GC integration. Theoretically, for an RNA molecule of length k, any of the four nucleotides can occupy each position, resulting in 4^k^ unique sequences (as illustrated in Fig. 1a). These sequences exhibit varying GC content, calculated as (G + C)/k * 100%, creating k+1 distinct GC-content categories. While GC content provides the percentage of G and C nucleotides, it doesn’t specify sequence or structure; thus, different sequences can share the same GC content. The distribution of RNA fragments within each GC-content category follows a Gaussian distribution (Eq. 4), derived from random permutation. The cumulative number of fragments across all GC-content categories is calculated using Eq. 5. By uniformly distributing individual RNA segments, we generate an idealized dataset mirroring the intact transcript template. This allows for accurate determination of key parameters, such as mean and standard deviation, for our self-benchmarking model. These predetermined parameters are then used to fit empirical datasets, mitigating biases and enabling accurate data standardization. Current bias correction methods struggle with uneven sequencing coverage arising from the complex interplay of known biases (GC bias, position bias, and sequence bias) and other, yet uncharacterized, biases. These methods typically derive single bias-weighting factors from empirical sequencing data, limiting their effectiveness. In contrast, GaussF takes a more effectiveapproach, extracting key structural parameters directly from the natural transcript sequence itself. By fitting the model’s cumulative function, parameterized with these pre-determined values, to the empirically observed, accumulated GC-categorized k-mer counts, we obtain bias-corrected transcript abundance estimates, effectively accounting for the complex biases present in the sequencing data.

**Fig. 1.**
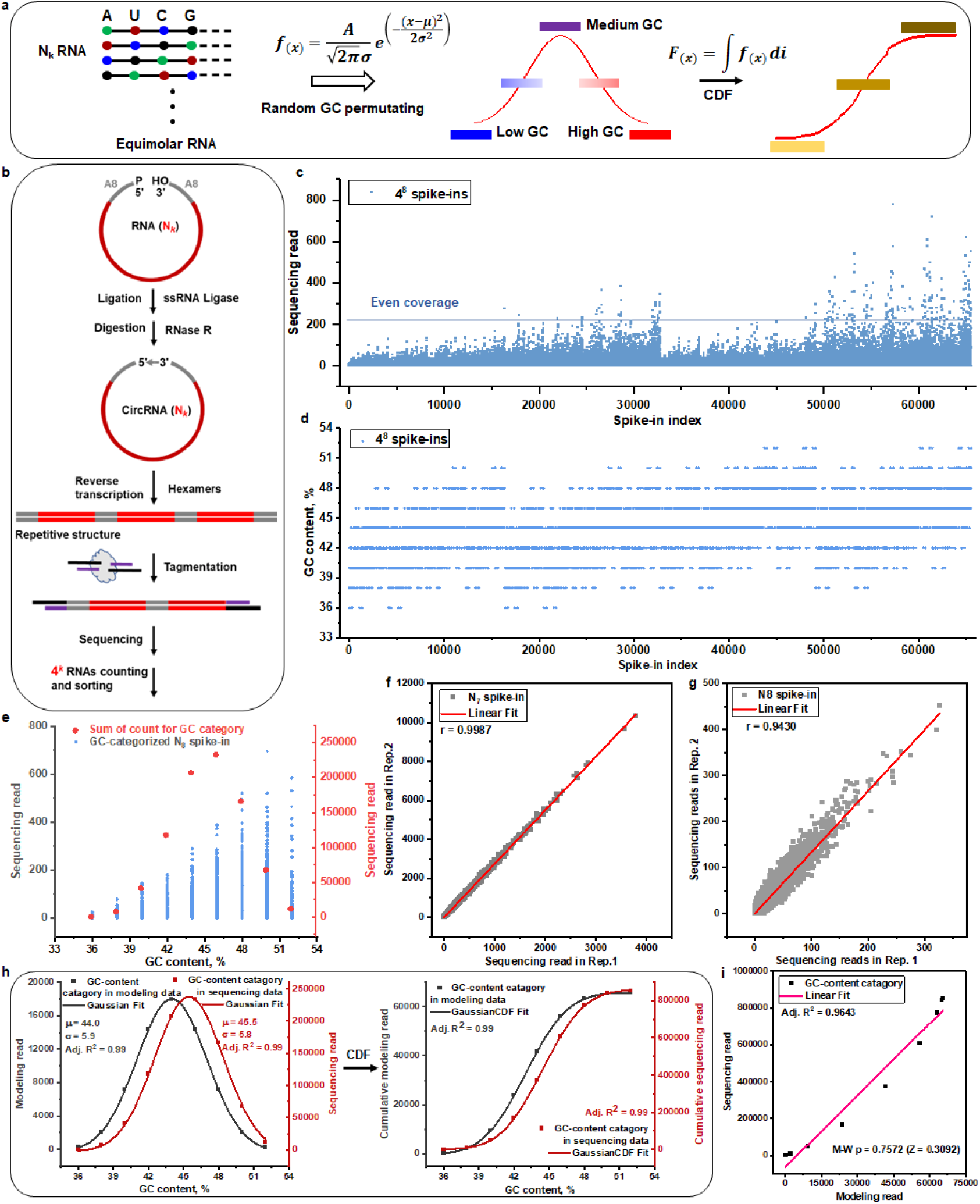
Parametric Gaussian model based on GC content. **a**) Conceptual foundations of GC-content quasi-Gaussian distribution function and a corresponding quasi-CDF. Random permutation of G and C bases gives rise to a Gaussian distribution for the GC content, mathematically derived from a binominal distribution. By utilizing this k-mer count Gaussian distribution function, a cumulative distribution function is inferred by integrating the k-mer count categorized by GC content. **b**) A sequential approach for validating GC content-dependent Gaussian distribution using RNA-seq. The RNA sequence with N randomly incorporated nucleotides, augmented with two identical A8 arms, undergoes ligation by single-strand RNA (ssRNA) ligase. Residual linear RNA is digested with RNase R. The remaining circular RNA templates encourage rolling circular reverse transcription, initiated by random hexamer priming. Sequencing adaptors are subsequently introduced via tagmentation and amplified by PCR. After sequencing, the data get analyzed based on cyclic consensus sequence. **c**) The sequencing read counts of each N8-incorporated (4^8^=65536) spike-in RNA (k-mer) are displayed for each alphabetically ordered nucleotide (A, T, C, G) at N positions. **d**) Presentation of GC-content values of each spike-in RNA, listed according to the alphabetical order of nucleotides at the N positions. **e**) The distribution of GC content-categorized read count. The sequencing read count corresponding to N8-incorporated spike-in RNA are grouped by the GC content. At each GC-content category, the sum of individual counts is presented, revealing a basic bell-shaped distribution. **f**) Correlation between N7 incorporated (4^7^=16384) spike-in RNA-seq replicates (n =2) is depicted. The linear fit produces a Pearson’s correlation coefficient of 0.9987. **g**) Correlation between N8 incorporated spike-in RNA-seq replicates (n=2) is displayed. A linear fit yields a Pearson’s correlation coefficient of 0.9590. **h**) The GC content-categorized read count distribution is presented as probability density or a CDF model, based on the sequencing reads and modeling reads. Parameters of the model, derived from sequencing and modeled reads, correspond to the Gaussian Distribution Function for OMF and a Gaussian function for cumulative distribution. **i**) The Probability-Probability (P-P) plot correlates the modeled data with the sequencing data for the cumulative read count based on GC content. The Kolmogorov-Smirnov (K-S) test yields a p-value of 1, attributed to the size of the sample, and the Mann-Whitney (M-W) test presents a p-value of 0.7572, based on the normalzied sequencing and modeling dataset within the 0-1 range. A linear fit is used for correction analysis.

In our quest to validate the self-benchmarking methodology and fully examine the inherent Gaussian distribution of GC content, circular RNA was employed as the subject for our RNA-seq evaluation (as depicted in Fig.1b). The sequencing process involved an initial phase of hexamer-centric reverse transcription, sparking tandem cDNA synthesis. This stage was followed by tagmentation and veiling through adapters, then came PCR amplification and, ultimately, sequencing. Including circular RNA in this investigation enhances the capacity to probe GC content, effectively steering the GC categorized-count distribution within the broad purview of RNA-seq. The process of hexamer-induced rolling circle amplification (RCA) generates cDNAs that extend beyond the size of the original circular RNA template.^29^ The RNA template is designed with a 50-mer core sequence interspersed with an 8-nucleotide random sequence (N8), and it is bordered by poly-A arms (8-mer) at both ends. These A8 arms, as illustrated in Fig.1b (top), were specifically engineered to counterbalance ligation bias during the assembly of equimolar circular RNA structures. We then embellished the synthesized cDNA products with adapters through the process of tagmentation, thus laying the groundwork for subsequent PCR amplification and sequencing. Throughout iterative sequences in data processing, the read count corresponding to each RNA sequence is determined (as shown in Fig.1c). Sequencing reads corresponding to each unique RNA sequence (N8 in a 50-mer) are systematically sorted using the alphabetical A, T, C, and G sequence. Interestingly, significant fluctuation is observed in the sequencing count, particularly when compared with evenly distributed coverage in modeling data. This phenomenon has been exemplified with nucleotide N7 in the context of a 25-mer spike-in RNA-seq experiment (as illustrated in supporting information Fig.S1, upper panel). This uneven coverage across all the spike-in RNAs can be attributed to various biases affiliated with RNA-seq. However, when sequencing counts for each spike-in RNA are sorted by GC content, a fairly stratified distribution emerges in nine (8+1) GC-content categories (as depicted in Fig.1d) and in eight (7+1) GC-content categories (as illustrated in supporting information Fig.S1, lower panel). Consequently, a Gaussian distribution of GC-categorized sequencing counts manifests regularly when individual counts linked to each GC-content category-enclosed RNA are accumulated (as illustrated in Fig.1e). Thus, the uneven distribution of sequencing counts across each spike-in RNA is transformed into a bell-shaped distribution of GC-classified counts, deeply rooted in the inherent GC permutation mechanism. Notably, the spike-in RNA with high and low GC-content are not underrepresented unpredictably but primarily due to inherent tendencies.

Further validating the reliability of our spike-in-based GaussF self-benchmarking model, we also incorporated a 25-mer spike-in RNAs along with N7 (as shown in supporting information Fig.S2), then subjected it for RNA sequencing paralleling the N8-50mer RNAs. Two iterations of spike-in-based RNA sequencing showed their matchmaking robustness through consistent correlations between replicates (as demonstrated in Fig. 1f and g). This alignment also laid the groundwork for comparing two essential data sets. The empirical uneven coverage characteristic of sequencing-generated data effectively corresponded with the theoretically even coverage of modelling-generated data across each GC-content category. Both sets, when arranged around the GC-categorized count, successfully fitted the Gaussian distribution function (as depicted in Fig. 1h). While analyzing the mean value divergence between the sequencing data and the modeling data, we noted a discrepancy of approximately 1.5 units in GC content within 16 GC-content intervals, hypothetically due to the limited sequence diversity predicated on only eight sites for random base loading. We anticipate that with a higher degree of GC-content diversity within naturally occurring transcriptomes, a well alignment would emerge. Despite this limitation, the standard deviations turned out to be closely matched in both data sets. The cumulative counts were in sync and their overlap for each category-enclosed RNA was effectively portrayed. Factoring in the shifted mean value, it was apparent that there were disparities positioned within the cumulative counting curves. Yet, both modeling and sequencing datasets aligned well with the cumulative distribution function. The P-P plot from data based on cumulated counts from both modeling and sequenced iterations supported the authenticity of this analogous cumulative distribution (as referenced in Fig. 1i). A strong correlation between datasets was found, as evidenced by a Mann-Whitney U test (M-W p) value of 0.7572. This compatibility ensures parameters from modeling data are viable for estimating transcript quantification based on sequencing data via parametric self-benchmarked GaussF processing.

### GaussF self-benchmarking model for characterizing human transcript

To confirm the reliability of the GaussF model, which uses GC content for self-benchmarking, rigorous evaluation of its precision and robustness is necessary, using experimental sequencing data from a natural transcriptome. Our methodology comprises the segmentation of a naturally-derived human transcript sequence from the 5’ end into specific sized bins, or k-mers, with the computation of GC content for each segment (as seen in Fig.2a). This approach is expected to shed light on the more intricate pattern of GC-content variability across the natural transcript, unlike the limited GC-content variation within the sequences generated via artificially N8-based methodologies. Nevertheless, the manifestation of GC-content complexity at the transcriptomic scale is intrinsically linked to the methods of library preparation adopted in the RNA-seq protocols. Here, we employ an RNA-seq library preparation protocol that adheres to a prevalent hexamer-priming scheme, a characteristic integral to the commercially available kits, including the VAHTS Universal V8 RNA-seq Library Prep Kit and the NEBNext® Ultra™ II RNA Library Prep Kit. This strategy ensures a consistent approach to hexamer targeting across all segments of the transcriptome, promoting theoretical uniformity in the representation of transcript regions. In contrast, oligo(dT)-anchored library preparation strategies target exclusively the poly(A) tails of mRNAs, a design element that introduces a pronounced 3’-end bias, resulting in a misrepresented range of GC content relative to the comprehensive diversity present in full-length transcripts. To illustrate the impact of these disparate library preparation approaches on read distribution patterns, comparative data from duplicated RNA-seq experiments conducted on HEK293T cells, using both hexamer-based and oligo(dT)-based protocols, are presented (as shown in Fig. 2b). The implementation of the hexamer-based RNA-seq methodology results in the distribution of mapped reads across the transcriptome landscape. Conversely, an oligo(dT)-centered protocol reveals the expected 3’ end concentration of reads, corroborating our hypothesis and illustrating the preferential targeting of the poly(A) regions within transcripts.

**Fig. 2.**
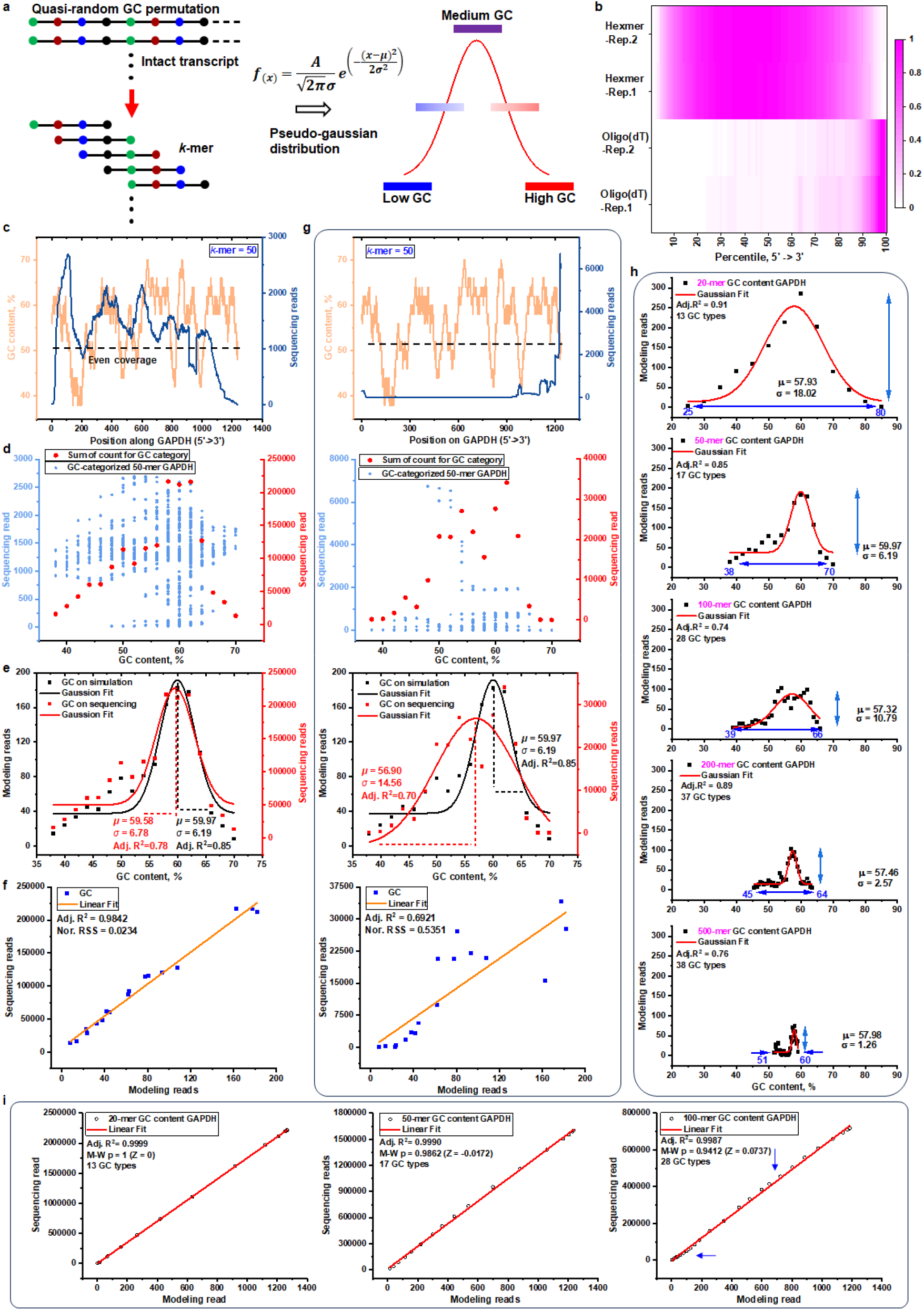
Assessment of GC content-based Gaussian distribution in natural transcripts. **a**) Derivation of a theoretical pseudo-Gaussian distribution for transcript GC content by integrating inherent sequence complexity, essential for formulating an accurate distribution representation. **b**) Comparison of read coverage obtained from hexamer RNA-seq and oligo(dT) RNA-seq across 59,100 transcripts, demonstrating the influence of library preparation on representation bias, analyzed using duplicates for each method (n= 2). **c**) Sequencing count and GC-content values of 50-mers along the 5’-start position of the GAPDH (ENST00000229239) human transcript are demonstrated, with evenly distributed 50-mers sequence coverage in the theoretical modeling data. **d**) Categorization and visualization of sequencing counts for GAPDH-derived 50-mers across 17 GC content classes, with aggregates signified by large spots to evaluate the GaussF model’s calibration capacity. **e**) Aggregated 50-mer counts depicted across GC content levels, overlaid with a pseudo-Gaussian distribution fitting curve, juxtaposing model parameters with sequencing metrics for comparative analysis. **f**) Linear regression assessing agreement between modeled and sequenced 50-mer count distributions, substantiated by an adjusted R² value of 0.9842 and measurement of data congruence through residual sum of squares (RSS), adjusted within the 0-1 range. **g**) GC content-specific distribution of GAPDH 50-mer counts in oligo(dT)-based RNA-seq. Distribution, compilation, and pseudo-Gaussian fit of GAPDH 50-mer sequencing and modeling counts, highlighting GC content-specific patterns and confirming model-read correlation with an adjusted R² of 0.6921. **h**) Analysis of k-mer size effect on GC content distribution within GAPDH sequences, indicating a narrowing range of distribution with increasing k-mer length. **i**) P-P plot comparing modeled and empirical cumulative read counts across different k-mer lengths, with significance tested by Mann-Whitney (M-W) based on the normalzied within the 0-1 range and linearity evaluated by linear fits, noting deviations from ideal correlation with directional arrows.

The GassF self-benchmark model’s effectiveness was verified through an in-depth examination involving hexamer-based and oligo(dT)-based datasets on a standard transcript, the glyceraldehyde-3-phosphate dehydrogenase (GAPDH). Sequences of 50-mers were extracted starting from the 5’ end of the GAPDH reference. The sequencing counts of each 50-mer, produced by the hexamer protocol, were plotted against the GC content of the corresponding segment (as depicted in Fig.2c). A theoretical model depicting even coverage across these segments was overlaid on the plot for comparative analysis. Contrary to the homogeneous distribution of 50-mers anticipated by the theoretical model, the empirical sequencing data unveiled intrinsic biases, resulting in an inconsistent coverage across the GAPDH sequence. The attempt to correlate 50-mer sequencing counts directly with their corresponding GC contents encountered analytical difficulties.^30, 31^ These disparities were further evidenced by the categorization of sequencing data by GC content, which delineated a distinctive and irregular pattern specific to the GAPDH transcript (as illustrated in Fig.2d, small dot). This deviation underscores the complexity inherent in translating raw sequencing data into a reliable representation of transcriptomic architecture. Within each GC content category, the occurrence of individual 50-mer sequences were aggregated (as illustrated in Fig.2d, big dot), producing an approximate bell-shaped distribution. This GC-categorized count distribution was then fit to a Gaussian distribution function (as depicted in Fig.2e). Concurrently, the distribution of 50-mer counts from the modeling data with even coverage was plotted, revealing close matches in fitting parameters between the hexamer-based sequencing data and modeling data. A strong linear correlation between the sequencing and modeling data further confirmed their accordance (as depicted in Fig.2f).

In the comparative analysis, oligo(dT)-based sequencing data presented a contrasting 50-mer distribution pattern that exhibited a pronounced preference for the 3’-end, deviating markedly from the even coverage observed across the GAPDH transcript (as depicted in Fig.2g). This pattern diverged significantly from that obtained from the hexamer-based dataset. Additionally, sequencing counts for each GC content category were aggregated and subjected to Gaussian distribution fitting. This analysis revealed substantial deviations in both mean and standard deviation compared to the projections of the theoretical model. This discrepancy was further confirmed by the correlation analysis of the GC content-categorized counts between the sequencing data and modeling data. The outcomes of our analysis suggest that the GassF self-benchmark model, designed to account for comprehensive GC content with an even coverage across entire transcripts in the modeling data, appears to be more suitable for normalizing datasets obtained from hexamer-based sequencing that targets full-length transcripts. The holistic scrutiny of transcripts’ GC content in hexamer-based datasets discloses a more exhaustive understanding of the GC content. In comparison, evaluations of GC content exhibit inherent limitations when conducted using oligo(dT)-based sequencing data, which inherently display a bias towards the 3’ termini of transcripts. However, it should be highlighted that the usage of oligo(dT)-based sequencing data can be well-standardized with an appropriately designed self-benchmark model capitalizing on evenly-distributed 50-mers, specifically within certain parts of the transcript’s 3’end region. This approach could potentially aid in maximizing the value acquired from oligo(dT)-based datasets. This way, each type of dataset can be adequately leveraged with their respective suitable self-benchmark models, highlighting the importance of selectivity when scrutinizing sequencing datasets.

Transcripts inherently possess unique sequences that generate distinct distribution profiles based on GC content. Nonetheless, our study found that the bin size, or the segment length used to calculate GC content, plays a crucial role in determining the distribution of read counts grouped by GC content. In our analysis of datasets modeled on GAPDH framework (as displayed in Fig. 2h), in addition to datasets derived from the ACTB and AMD1 genes (as shown in the supporting information Fig. S3), an increase in fragment length led to a consequent fall in goodness of fit. Specifically, when the bin size for GC content calculation extended from 20 to 500 bases, there was a discernible decrease in the range of both GC content and its corresponding read count. This reduction is probably due to the larger bin size capturing a wider array of GC content values which, despite their diversity, tend to converge towards the mean. This densification, in turn, caused the orderly pattern of the initial GC content-based distribution to become more amorphous. This phenomenon could elucidate why read counts allocated to GC content in standard GC-bias models are highly vulnerable to external influences within the sequencing pipeline.^32^ To validate these observations, the dataset was subjected to analysis with predetermined lengths-20-mer, 50-mer, and 100-mer-within the GaussF model framework (as delineated in the supporting information Fig. S4). Thereafter, we assessed the distribution patterns of read counts stratified by GC content, drawing a comparative analysis between computational modeling data and empirical sequencing data derived from the GAPDH transcript (as depicted in Fig. 2i). Intriguingly, we noticed an upward deviation from the diagonal line signifying a leftward skew of the observed cumulative distribution compared to the theoretical distribution. This skew suggests a higher occurrence of lower-valued outcomes in the dataset. Furthermore, a portion of the curve fell below the theoretical linear line indicating a rightward skew of the observed cumulative distribution relative to the theoretical distribution, demonstrating a surplus of higher-valued results in the sample. The skewed tendency is not only evident within the high-abundance transcripts, such as those of GAPDH, but also becomes more pronounced in transcripts exhibiting lower abundance. For instance, a distinct skewness is manifest in the dataset corresponding to the transcript EMP1-211, as evidenced by the notable deviations in both upward and downward directions from the diagonal line (as shown in supporting information Fig. S5). Significantly, as the k-mer length increased, the goodness of fit in terms of data normalization accuracy diminished. As such, careful consideration should be given to determining an optimal k-mer length to ensure accurate normalization of transcript data.

### Workflow for Isoform Alignment and Fitting Procedure

To prepare for k-mer counting from sequencing data, we first generate transcript-specific k-mer reference sets using the reference isoform sequences (e.g., obtained from Ensembl). This process involves systematically segmenting each individual isoform sequence (as shown in Fig. 3a). Specifically, starting from the 5’ end, each transcript is divided into overlapping segments of a predefined length, k (e.g., 50 nucleotides), covering its entire length. This procedure ensures that a comprehensive collection of all possible k-mers originating from that specific isoform is created. Repeating this for all isoforms yields distinct k-mer sets unique to each transcript variant. These sets serve as the reference dictionary against which k-mers detected in sequencing reads will be quantified in downstream analysis. Notably, when analyzed, the collective GC content distribution across these k-mer sets derived from natural transcript sequences displays a diversity comparable to that observed in sequences synthesized through random base incorporation (A, U, C, G).

**Fig. 3.**
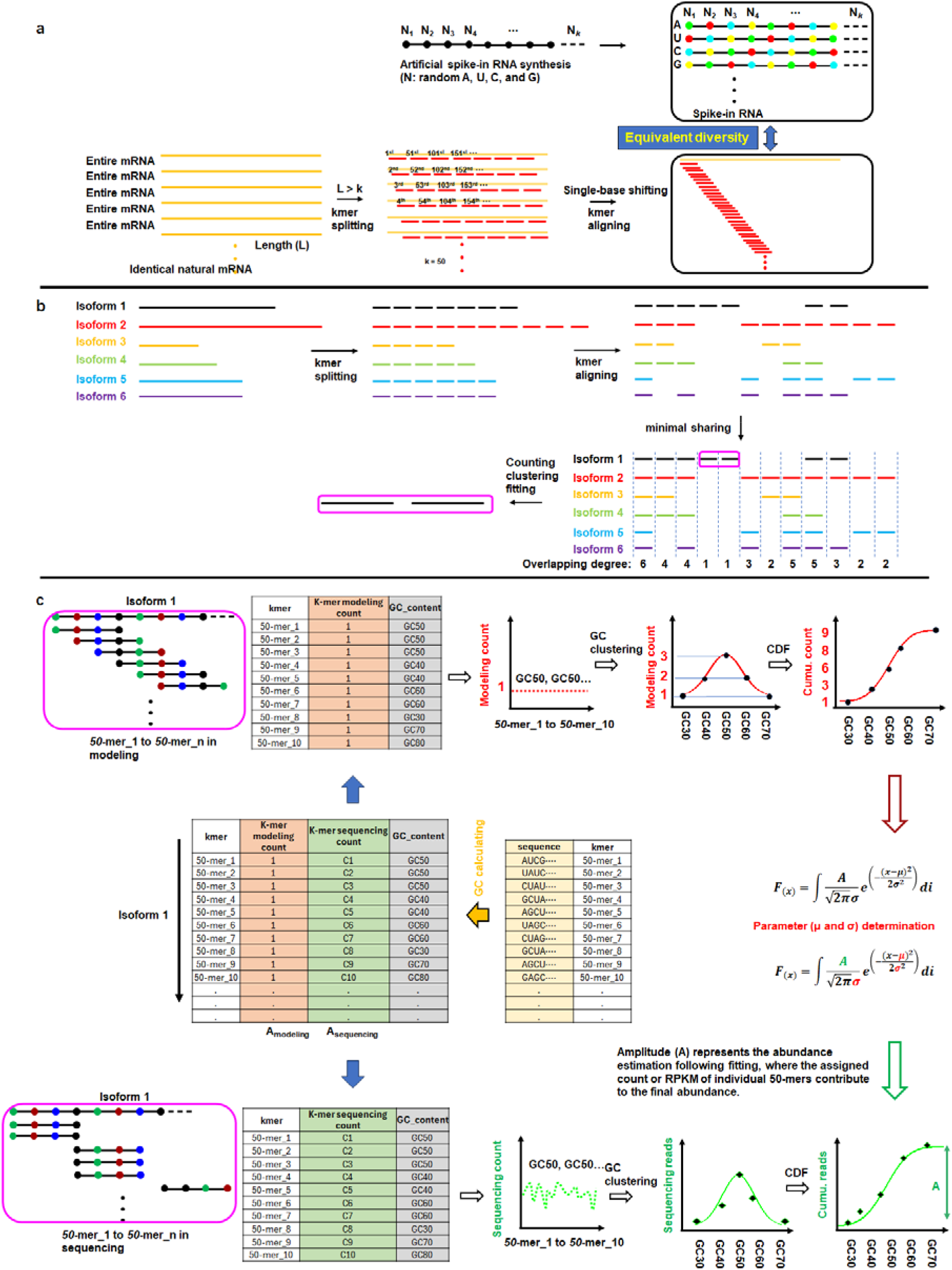
Parametric Gaussian CDF-based self-benchmark model for transcript quantification. **a**) Spike-in RNAs were synthesized with random incorporation of A, U, C, and G. Transcript sequences for relevant species (e.g., Human: GRCh38.cdna.all.fa) were downloaded from Ensembl. Each transcript was computationally split into overlapping k-mers starting from the 5’ end to define its specific k-mer set. **b**) All isoform transcript sequences (Homo_sapiens.GRCh38.cdna.all.fa) were downloaded from www.ensembl.org. Each isoform was split into kmers starting from the 5’ end. These isoform-specific kmer sets were then aligned across the entire transcriptome kmer pool to calculate the global frequency in terms of overlapping degree. Based on this overlapping degree, the minimal shared regions (kmers) were identified for downstream GC-based clustering, counting, and fitting. **c**) Transcript-specific benchmarking parameters are ascertained by categorizing evenly-distributed 50-mers throughout the transcript based on their GC content, and integrating individual occurrence counts within each GC category. The table show the data acquisition from the sequence data and transcript sequence. The columns K-mer modeling count and GC content are used to determine the key parameters. The resultant GC content-enumerated counts are then fit to a pseudo-OMF, and an accumulative distribution function (CDF) is applied to accommodate the cumulative GC content-related counts. This procures the two fundamental parameters: mean (μ) and standard deviation (σ). Uneven k-mer coverage exhibiting position-specific occurrence counting facets of the sequencing data form the columns of kmer-sequencing count (or normalized kmer sequencing count) and associated GC content. The k-mers, categorized by GC content, are summed up within each category, and plotted. Once the parameters derived from the modeling data are set in place, the pseudo-OMF aptly acknowledges all intrinsic biases in the sequencing data. Additionally, by applying the pseudo-CDF with the fixed parameters, accurate estimates of transcript abundance are achieved by taking into consideration the cumulative effect on k-mer occurrence count.

The isoform alignment workflow first involves segmenting each isoform sequence into k-mers of a defined length (e.g., 50 nucleotides), using either overlapping or non-overlapping windows. All resulting k-mers from the entire transcriptome are then collected into a comprehensive set. These k-mers are subsequently aligned against this set (as illustrated in Fig. 3b) to identify matches. This alignment serves two primary purposes: first, to identify k-mers shared between different isoforms of the same gene, which helps map common regions; and second, to find potential matches between k-mers originating from different genes, allowing exploration of broader sequence similarities or motifs. Following this global alignment, the analysis narrows its focus to each specific gene. Within a gene, the minimal set of k-mers shared across its various isoforms is determined, highlighting conserved sequences. Concurrently, unique k-mers characteristic of particular isoforms are identified to enable their distinction. Ultimately, these sets of minimal shared and unique k-mers allow for the definition and characterization of isoform-specific regions, which may correlate with functional domains.

Building on the alignment results, the fitting procedure commences by preparing an initial data table for each k-mer located within the identified minimal shared transcript regions. As shown in Fig. 3c, this table consolidates key information for each k-mer: its sequence, an initial modeling count (arbitrarily set to 1), its calculated GC content, and the empirical read count derived from processed sequencing data. The empirical counts may incorporate normalization metrics such as RPKM (Reads Per Kilobase per Million mapped reads) or TPM (Transcripts Per Million). Using this data structure (illustrated with a representative set), k-mers are grouped based on identical GC content values. A Gaussian distribution, reflecting the expected patterns of GC content in genomic sequences, is then fitted to the distribution of these GC content groups. From this fitted Gaussian model, the Cumulative Distribution Function (CDF) is derived. Critically, the parameters essential for the subsequent quantification model-namely the mean and standard deviation characterizing the GC content distribution-are estimated directly from this theoretical Gaussian CDF. This parameter estimation depends solely on the GC composition distribution inherent in the modeled k-mers and is performed independently of their associated empirical sequencing read counts at this stage.

### Transcript-specific GaussF model for unbiased quantification

The self-benchmarking GaussF model presents a significant advancement in addressing complex biases encountered within sequencing data. It employs a OMF and a corresponding pseudo-CDF for the meticulous determination of key parameters required for data parametric calibration. This dual approach allows for a theoretically uniform coverage of k-mers throughout the transcriptome, facilitating the processing of unpredictably bias-imbued sequencing datasets. In particular, the GaussF model establishes a transcript-specific benchmark by leveraging two principal parameters: mean and standard deviation, based on the even coverage of k-mers. These parameters, which are clearly delineated in Fig. 4a, serve as the foundation for the pseudo-Gaussian distribution function, enabling it to competently mitigate all biases intrinsic to sequencing data. The pseudo-CDF further capitalizes on these parameters by aggregating the cumulative effect on k-mer counts, thereby rendering the unbiased estimates of transcript abundance (as demonstrated in Fig. 4b). This methodical, transcript-oriented benchmarking framework potentially excels beyond traditional empirical approaches, providing refined transcript quantification. A step-by-step procedural description (as shown in Fig. 3) is able to delineate the computational procedure more clearly. The ideal of achieving even k-mer coverage across the full-length transcript represents an aspirational benchmark for bias-free datasets, which current quantification methodologies continually strive to attain through various algorithmic means, including the Expectation-Maximization (EM) algorithm.^17, 20^ While the EM algorithm has been prominent due to its computational efficiency in managing the convergence of effects arising from multiple biases, the application of the EM algorithm may be constrained by the complexity and diversity present in sequencing read profiles. Attaining an outcome analogous to bias-free, even coverage through the EM method presents notable difficulties. The GaussF model embraces a self-benchmarking strategy that aligns with the concept of even theoretical coverage. By implementing this refined framework, the GaussF model endeavors to comprehensively counteract bias, with the objective of enhancing the accuracy and reliability of transcript abundance measurements.

**Fig. 4.**
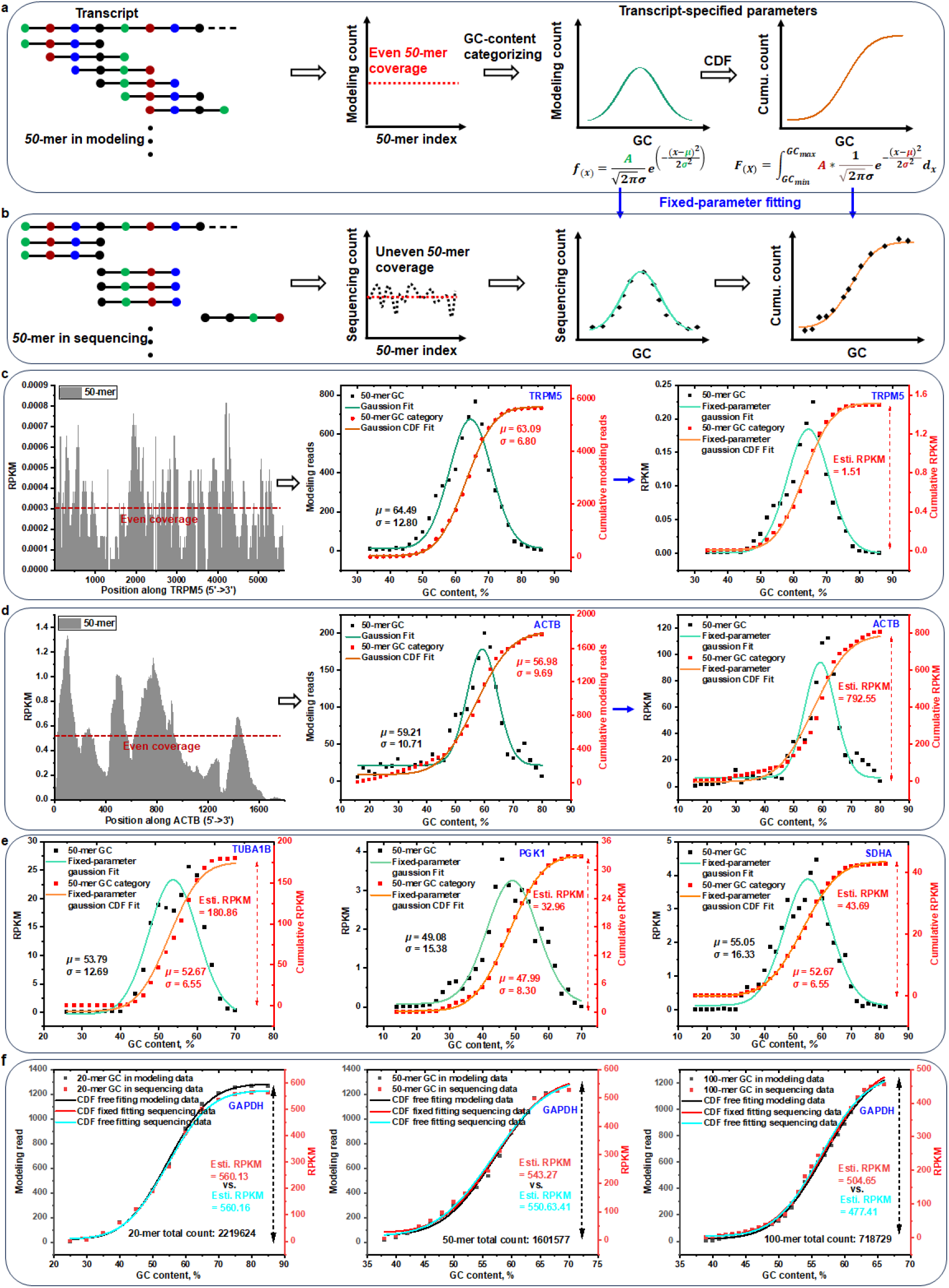
Parametric Gaussian CDF-based benchmark model for transcript quantification. **a**) Transcript-specific benchmarking parameters are ascertained by categorizing evenly-distributed 50-mers throughout the transcript based on their GC content, and integrating individual occurrence counts within each GC category. The resultant GC content-enumerated counts are then fit to a pseudo-OMF, and an accumulative distribution function (CDF) is applied to accommodate the cumulative GC content-related counts. This procures the two fundamental parameters: mean (μ) and standard deviation (σ). **b**) Uneven k-mer coverage exhibiting position-specific occurrence counting facets of the sequencing data. The k-mers, categorized by GC content, are summed up within each category, and plotted. Once the parameters derived from the modeling data are set in place, the pseudo-OMF aptly acknowledges all intrinsic biases in the sequencing data. Additionally, by applying the pseudo-CDF with the fixed parameters, accurate estimates of transcript abundance are achieved by taking into consideration the cumulative effect on k-mer occurrence count. **c**) The pseudo-CDF-based model is applied to quantify transcript TRPM5 (ENST00000696290). Theoretical even coverage are overlapped with empirical uneven coverage to pinpoint parameters such as benchmarking mean (μ) and standard deviation (σ). The parameter-fixed pseudo-OMF alongside the pseudo-CDF is then used for normalizing the biased data and estimating cumulative RPKM. Individual RPKM is allocated to each 50-mer (k-mer) present along the transcript. **d**) Application of the pseudo-CDF-based self-benchmarking model for the quantification of ACTB transcript (ENST00000646664). **e**) The self-benchmarking quantification is shown for the TUBA1B (ENST00000336023), PGK1 (ENST00000373316), and SDHA (ENST00000264932) transcripts with different transcript sizes and intrinsic expression levels. **f**) Examination of the k-mer effect on transcript quantification, concentrating on the GAPDH transcript (ENST00000229239). Discerned parameters for 20-mer, 50-mer, and 100-mer based functions applied to RPKM values are fitted by two distinct modeling approaches: a parameter-fixed model and a parameter-free model.

In an attempt to validate the theoretical self-benchmarking process, we performed the task of quantifying transcript abundance from human tissue by going through various stages of data processing. This process involved steps such as the tallying of k-mer sequences and their initial normalization, stratification of GC content, ascertainment of relevant parameters, and employments of fixed-parameter fitting techniques (as depicted in Fig.4c). The counts (as shown in supporting information Fig. S6) for each k-mer along transcript were transformed into standard RPKM for analysis. Given that the majority of processed sequencing reads are composed of 150 nucleotides (as depicted in Supplementary Fig. S7), it can be inferred that a single read has the theoretical potential to generate 101 distinct 50-mer sequences under an optimal base-calling framework. This calculation is predicated on the sliding-window approach wherein each consecutive 50-nucleotide segment within the 150-nucleotide read is considered a unique k-mer. RPKM was therefore expressed through the formula RPKM = countk*2*1000*1000000/(101*lengthT*readT) or TMP was also indicated by the formula TPM = countk*2*1000*1000000/(101*lengthT*sum_RPK), where countk symbolizes the sequencing count of all k-mers from the transcript, lengthT refers to the entire transcript or minimal shared-region length, readT signifies the total read count in the fastq data and sum_RPK means the sum of all individual RPK (Reads Per Kilobase) values that is calculated as the number of reads mapped to this transcript divided by its length in kilobases calculated for every transcript in the dataset. This normalization process for isoform-level transcript quantification utilizes the length of the minimal-shared region specific to each transcript to ensure accurate representation of transcript abundance. We graphed the RPKM of each 50-mer along TRPM5 (as depicted in Fig.4c, left). Basing our data on the equal coverage of TRPM5 in the model, we evenly distributed a preset of 5640 50-mers for the categorization of GC content, before summing them to fit the Gaussian distribution and the pseudo-CDF. Key parameters including the mean and standard deviation were then determined and utilized to amend the GC content-categorized 50-mer sequencing count distribution (as shown in Fig.4c, middle). The elimination of inherent biases was achieved through the fitting of the Gaussian distribution, and the amount of TRPM5 estimated through cumulative fitting (as shown in Fig.4c, right). The self-benchmarking methodology was further applied to estimate the abundance of a ubiquitously expressed reference transcript, ACTB (as shown in Fig.4d). Consistent with our expectations, the estimated abundance levels of ACTB markedly exceeded those of TRPM5, corroborating the robustness of the approach. To underscore the adaptability of the GaussF model, we expanded our analysis to include transcripts characterized by varied structural complexities and intrinsic abundance levels, namely TUBA1B (1627 nucleotides), PGK1 (4812 nucleotides), and SDHA (2693 nucleotides) (as shown in Fig.4e). These assessments allowed us to demonstrate that the GaussF model adeptly reconciles the self-benchmarking parameters to adjust for biases, thereby facilitating accurate and reliable quantification of transcript abundance for a diverse array of transcripts.

To investigate the influence of k-mer size on the precision of transcript quantification, we assessed the abundance of the GAPDH transcript by analyzing k-mers of differing lengths, encompassing 20 nucleotides (nt), 50 nt, and 100 nt (as shown in supporting information Fig. S8). Our analysis was anchored on the application of a cumulative function with a fixed mean and standard deviation to fit the cumulative sum sequencing count of GC content sorted by k-mers lengths. Each k-mer length delivered unique parameters that informed the cumulative count distribution based on our model’s even coverage data. It became clear that the fitting abundance declined consistently with the increase of k-mer length. This trend could be attributed to the heightened sensitivity of longer k-mers to base-calling errors appearing in the reads. Upon comparing the abundances estimated using fixed and free-fitting models for varying lengths of k-mers (as depicted in Fig.4f), we noted an escalating disparity as k-mer length increased-in stark contrast to the 20-mer case where the abundance derived from both models aligned closely. This divergence in estimations underscores the effectiveness of the GaussF model in empirically normalizing data. However, it is noteworthy that shorter k-mers, such as the 20-mer set, can only generate a limited number of GC content types. Within the full length of the GAPDH transcript, a mere 13 distinct GC content categories are discernible when employing a 20-mer k-mer configuration. For transcripts of notably short length, the assortment of potential GC content categories is further diminished. This paucity of diversity may be inadequate to encompass the full spectrum of structural variation present within the data, consequently leading to an unrepresentative quantification of the population stratified by GC content. This could, in turn, lead to powerless normalization using preset parameters. Moreover, ensuring uniqueness for specific transcripts can become quite challenging due to the short k-mer set during k-mer counting. In light of these observations and challenges, we determined that a 50-mer length is optimal for subsequent abundance estimation via our parametric GaussF model.

To comprehensively demonstrate the effectiveness of GaussF, we analyzed RNA-seq data simulated at a transcriptome-wide scale, where the ground truth abundance for each isoform transcript was predefined for tracking purposes. We utilized Polyester Simulation to generate this RNA-seq data.^33^ It is crucial to acknowledge that the statistical distribution of fragment lengths-whether normal or otherwise-employed by Polyester or any other simulation tool, significantly influences which regions of the transcript are sampled. As a result, fragments are not uniformly distributed along the transcript length, leading to non-uniform coverage. Crucially, for isoforms with higher expression levels, a larger number of fragments is generated. However, this does not necessarily mean that each fragment within those isoforms has an equal probability of being selected. This non-uniform sampling introduces biases in the dataset, affecting the coverage profile of each isoform. We confirmed this issue using another simulator, which also produced data with significant non-uniform coverage, particularly noticeable drop-offs at the 3’ and 5’ transcript termini. This indicates an inherent bias in current simulation approaches. The implications of this skewed sampling are profound when interpreting the ground truth abundance of isoforms in RNA-seq data. The presence of an isoform in the dataset does not ensure equal representation of all its segments. This inherent bias in the data affects coverage profiles, which deviates significantly from an ideal scenario where coverage across fragment distributions would be even (as illustrated in supporting information Fig. S9a). These considerations highlight the importance of being cautious in interpreting simulated RNA-seq data, particularly when analyzing and comparing the ground truth abundance of isoforms. The biases introduced by non-uniform fragment sampling must be taken into account to accurately assess the effectiveness of analytical tools like GaussF in realistically complex scenarios. To facilitate a controlled comparison under simplified conditions, we evaluated estimates from GaussF, Salmon, Cufflinks, and Kallisto against ground truth normalized abundances using specifically designed simulated data (as shown in supporting information Fig. S9b). In this simulation, we focused on 100 target transcripts, including all ACTB and GAPDH isoforms. These were assigned an equal and relatively high abundance (preset relative value = 20). This equal abundance setting allows for a more straightforward comparison of method-specific artifacts, removing the complexity of differential expression. All other transcripts were included at an equal, low abundance to serve as background. We limited the set of highly expressed transcripts to 100 primarily due to computational constraints: simulating the high read coverage across all isoforms would generate prohibitively large FASTQ files and significantly increase processing time. For these 100 transcripts, each assigned a preset relative abundance of 20 in Polyester, we calculated the ground truth normalized abundance. This was determined using the formula: (Preset Relative Abundance * Scaling Factor * 1000) / Transcript Length, which translates to (20 * 9.66 * 1000) / Transcript Length in this case. This calculation incorporates Polyester’s internal scaling factor (≈ 9.66) needed to reach the target read count, and normalizes by transcript length (scaled by 1000) to produce a value analogous to RPK. We then assessed tool performance by correlating their abundance estimates with these ground truth values (Fig. S9b). Salmon (r = 0.83), Cufflinks (r = 0.82), and Kallisto (r = 0.81) achieved high Pearson correlations, whereas GaussF showed a lower correlation (r = 0.64). GaussF, focusing on minimal shared transcript regions, deviates in correlation analysis, as it may miss or underestimate transcripts with no reads in their unique regions. For example, GAPDH_ENST00000619601 shows null abundance with GaussF, where no reads or k-mers were detected, while other methods reported detectable values (as shown in supporting information Fig. S9c). The unique approach of GaussF, focusing on minimal shared regions, can lead to underestimation or complete omission of particular isoforms where the unique regions lack sufficient sequencing evidence. This outcome underscores the variations in correlation analysis and highlights the critical need for considering method-specific constraints and assumptions in the quantification of isoforms using RNA-seq data.

### Transcript isoform-level quantification with GaussF model

The GaussF method has been extended to achieve isoform-level resolution in RNA sequencing. Isoform-level quantification is particularly challenging because genes often produce multiple isoforms via alternative splicing, many of which share identical sequences. In such cases, the 50-mer coverage at each position may represent a mixture of splice variants rather than a single isoform (see supporting information Fig. S10). Moreover, every full-length transcript is composed of multiple distinct regions that may recur in various isoforms, complicating the accurate assignment of sequencing reads. This challenge is exemplified by our analysis of the EMP1 gene, which generates twelve annotated isoforms (see supporting information Fig. S11). To address these challenges, we developed a “minimal-shared” strategy that segments the full-length transcript into specific regions based on their association with different isoform configurations. These regions fall into two main categories: regions unique to a single isoform and regions shared by multiple isoforms (e.g., bi-isoform, tri-isoform, or more). The unique regions serve as definitive markers for individual isoforms, while the minimal shared regions delineate overlapping groups by identifying the smallest common sequence blocks. Such minimal shared-sequence regions for single isoform typically align with alternative splicing sites, particularly in less abundant (minor) isoforms,^34^ or within the predominant isoform, allowing for straight-forward assignment to corresponding isoforms. This strategy effectively simplifies the otherwise complex isoform landscape; for example, among nine EMP1 isoforms, we were able to identify unique regions specific to six isoforms (EMP1-201, EMP1-204, EMP1-205, EMP1-208, EMP1-210, and EMP1-211) (see Fig. 5a). To ensure unbiased quantification, the GaussF model was incorporated using counts of unique 50-mer sequences within each isoform. The cumulative GC content-categorized RPKM calculated from these unique regions for the six EMP1 isoforms was consistent between both parameter-fixed and free models (as illustrated in Fig.5b). With even coverage, the empirical GC-categorized RPKM can be normalized to theoretical parameters, enabling an unbiased estimation of transcript abundance. Furthermore, unique regions sometimes display distinctive patterns of sequencing coverage. For instance, four isoforms (EMP1-202, EMP1-203, EMP1-206, and EMP1-209) show no detectable 50-mer counts within their unique regions (as represented in Fig.5c). This absence creates an observable dip in the full-length 50-mer count profile corresponding to the minimal shared region, marked by an abrupt drop in counts when transitioning from the common to the unique regions, followed by a sharp increase when returning to the common region. While RNA-seq biases typically lead to gradual fluctuations in k-mer counts, these acute changes suggest that the unique regions more accurately reflect the true expression levels of their respective full-length isoforms. In these instances, the proportionality of unique region counts to overall isoform abundance is a reliable proxy for quantification. However, for these particular four isoforms, any misassignment of counts from the shared regions could significantly skew the estimates. Therefore, to maintain accuracy, we exclusively used counts from the unique regions when applying the GaussF model for isoform quantification.

**Fig. 5.**
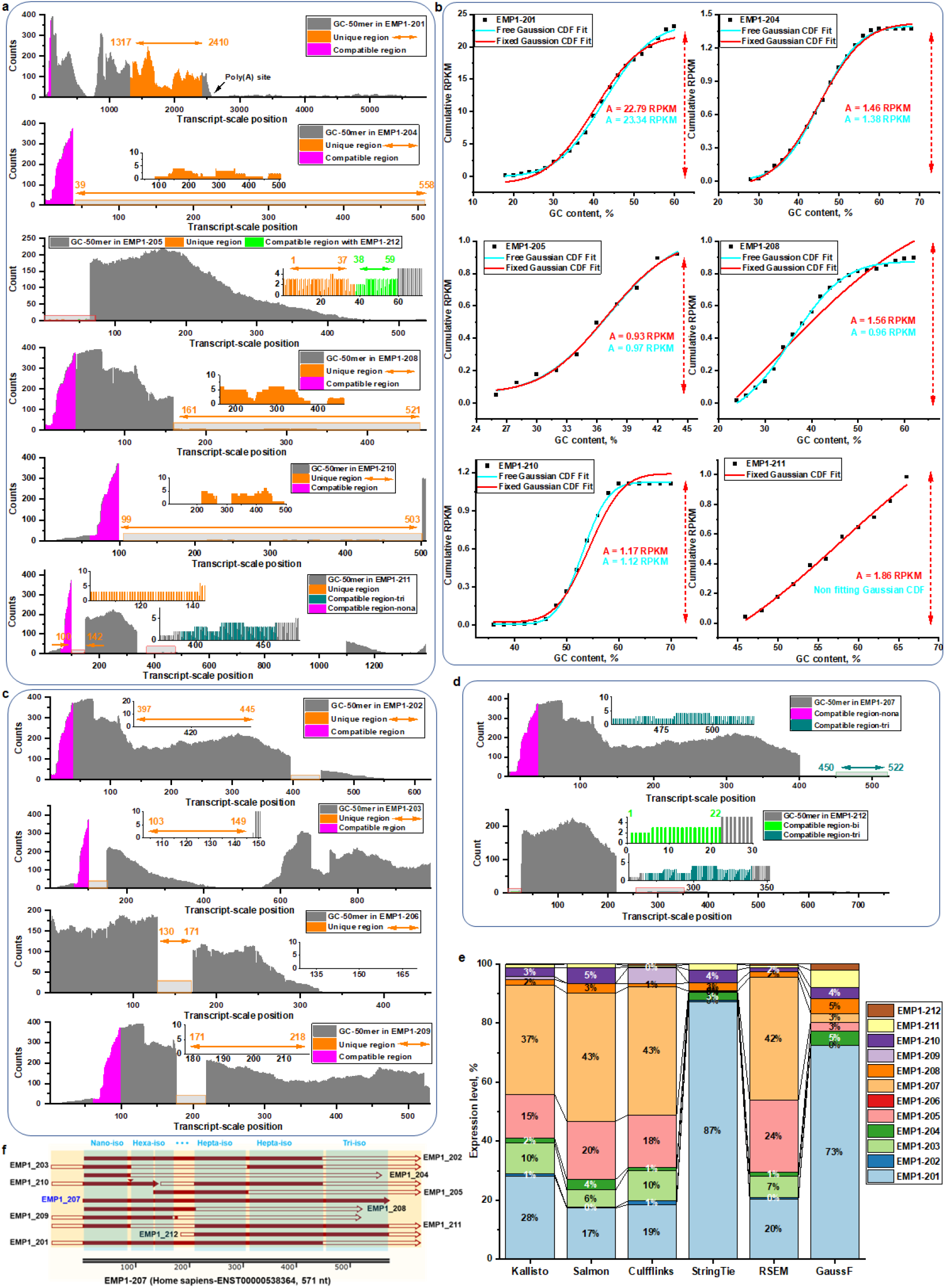
Isoform-level transcript quantification enabled by the GaussF model with self-benchmarking capabilities. **a**) Profiling and parametric quantification of six EMP1 isoforms. Detailed exploration of six EMP1 gene-derived isoforms-EMP1-201 (ENST00000256951), EMP1-204 (ENST00000535134), EMP1-205 (ENST00000536383), EMP1-208 (ENST00000541935), EMP1-210 (ENST00000542474), and EMP1-211 (ENST00000544053)-using 50-mer sequencing counts for transcript abundance profiling is depicted. Quantification analysis utilizes a parametric pseudo-CDF focused on the 50-mer counts along unique regions of the isoforms. A nona-isoform shared region at the 5’-start position, common across these variants, demonstrates a uniform pattern of expression within this conserved segment. The inset provides a magnified view, delineating the unique regions attributed to each isoform and thereby illustrating the differential transcriptional architecture. **b**) Illustration of the self-benchmarking approach for quantifying isoforms. GaussF model is applied to cumulative 50-mer sequencing counts that are normalized to RPKM. Parameter-fixed and free models provide both unbiased and biased evaluations of isoform abundance. **c**) Undetectable isoforms, EMP1-202 (ENST00000396301), EMP1-203 (ENST00000431267), EMP1-206 (ENST00000537612), and EMP1-209 (ENST00000542289), are presented. Each full-length isoform is profiled using a 50-mer count, and a nona-isoform shared region is highlighted. Unique regions within the profile are indicated by pairs of arrow lines. **d**) Display of the isoforms EMP1-207 (ENST00000536383) and EMP1-212 (ENST00000546357) with a tri-isoform and bi-isoform shared region, detailing their profiles based on the 50-mer count. The EMP1-207 isoform is profiled with 50-mer sequencing count, showing a tri-isoform shared region with minimal isoforms, EMP1-211 and EMP1-212. A nona-isoform shared region is also highlighted. The EMP1-212 shared a region with EMP1-205. **e**) A stacked bar chart presents the abundance proportions of the twelve EMP1 isoforms, determined by various transcript quantification methods such as Kallisto, Salmon, Cufflinks, StringTie, RSEM, and GaussF. The chart portrays the relative contributions of each isoform to the total mapped reads across all methods. **f**) Alignment of all EMP1 isoforms with reference to EMP1-207 is performed, spotlighting nano-iso, hexa-iso, hepta-iso, and tri-iso (isoform) shared regions.

In a typical scenario, the sequencing read count or RPKM for a transcript is derived from all reads mapped to both unique isoform-specific regions and regions common among multiple isoforms. Traditional quantification methods attempt to capture all mapped reads from the full-length transcript to estimate abundance. Reads from unique regions are straightforward to assign, but those from shared regions pose a significant analytical challenge. Variable count distributions in different shared regions can introduce bias, especially when evaluating low-abundance isoforms; even minor redistributions of counts may substantially alter the final assessment. To address this issue, we developed an approach that categorizes regions based on the degree of overlap among isoforms. When an isoform lacks a unique region, there may be another region that overlaps with one or more other isoforms-this shared area then represents the isoform for 50-mer counting. If neither a unique region nor a bi-isoform overlapping region exists, the simplest representative area might encompass three isoforms, resulting in a tri-isoform overlapping region. This concept can be extended further to tetra-or penta-isoform overlaps as needed. Essentially, this strategy allows us to pinpoint the simplest common region that collectively represents specific isoforms. For example, a bi-isoform region is identified in EMP1-212 (as shown in Fig. 5d). In this case, the emphasized region in EMP1-212 is common with EMP1-205, even though it differs from the unique region associated with EMP1-205. Consequently, using the bi-isoform region as a basis, the GaussF model processes the 50-mer counts for RPKM estimation for EMP1-212, with the resulting abundance reflecting a collaborative contribution of both EMP1-205 and EMP1-212 isoforms. Similarly, a triple-isoform region is identified for EMP1-207. This region is shared among EMP1-207, EMP1-211, and EMP1-212 and is used for abundance estimation of EMP1-207, as EMP1-211 and EMP1-212 have their own distinct unique regions. Moreover, this triple-isoform region does not overlap with the unique regions specific to EMP1-211 or EMP1-212. Thus, the abundance measured from the triple-isoform region reflects the combined contribution from these three isoforms.

To substantiate the precision of the unique region-based GaussF model, we undertook a comparative analysis with several typical isoform-level quantification approaches such as Kallisto, Salmon, Cufflinks, StringTie, and RSEM. These approaches are anchored firmly on alignment-based or alignment-free strategies. With regard to a set of identical sequencing data, these methodologies were used to quantify the twelve EMP1 isoforms (as illustrated in Fig.5e). The stacked bar chart graphically displays the relative abundance of each transcript isoform, normalized to reflect their proportionate contribution to the total expression. This visualization facilitates the comparison of isoform expression levels within the sample, while the absolute expression values (as shown in supporting information Fig. S12) are concurrently presented to provide quantitative context to the isoform-specific expression data. Interestingly, the expression profiling methodologies employed by Kallisto, Salmon, Cufflinks, and RSEM exhibit congruent distribution patterns, with EMP1-207 consistently emerging as the most abundantly expressed isoform among its counterparts. As previously identified, there is no unique region for EMP1-207, but several distinct regions shared with other isoforms (as shown in Fig.5f). The ‘triple-isoform’ region, shared by exactly three isoforms, serves as the simplest representative region for EMP1- 207-unlike other regions that are common to more than three isoforms (see supporting information Fig. S11). Within this triple-isoform region, the 50-mer counts are allocated to all three contributing isoforms. Notwithstanding, this region displays a substantially diminished level of 50- mer counts in comparison to other regions, which introduces a confounding discrepancy when contrasted with the predominance of EMP1-207 as indicated by quantification tools such as Kallisto, Salmon, Cufflinks, and RSEM. We propose that this apparent high abundance of EMP1- 207 is likely an artifact, resulting from the misattribution of k-mer counts or mapped reads from complex regions shared across multiple isoforms. To investigate this, we first established a baseline using EMP1-201. This isoform features unique sequence regions with high k-mer counts, facilitating reliable quantification. Consistently, all evaluated methods, including our approach GaussF, assigned high abundance to EMP1-201, confirming GaussF’s accuracy for less ambiguous targets. In contrast, EMP1-207 completely lacks unique regions; its sequence is composed entirely of segments shared with at least two other isoforms (minimum shared degree of 3, see Fig. 5d). GaussF utilizes k-mer counts specifically from these minimally shared regions. Our analysis found that the combined signal originating from all isoforms sharing these regions was relatively low. Therefore, since EMP1-207’s individual contribution cannot logically exceed this low shared signal, GaussF accordingly assigned it a low abundance. Intriguingly, although no counts are detected in the unique region of EMP1-203 (referred to Fig.5c), its expression is still identified by methods such as Kallisto, Salmon, Cufflinks, and RSEM. This discrepancy likely arises because these tools rely on EM algorithms for assigning ambiguous reads, enabling them to detect isoforms even when unique region counts are absent. In contrast, the StringTie approach applies a stringent classification process to ambiguous reads by leveraging precise, single alternative splicing junction graphs for differentiating isoforms.^16^ As a result, StringTie does not provide an abundance estimate for EMP1-203, which also helps avoid the overestimation of EMP1-207’s expression. These findings align with the outcomes derived from the GaussF model, suggesting that methodologies with meticulous read assignment strategies tend to yield more conservative estimates of isoform expression levels. However, it is important to note that under different assembly conditions, errors in isoform expression detection can still occur.

Glyceraldehyde 3-phosphate dehydrogenase (GAPDH) is frequently utilized as a housekeeping gene in human sample analyses due to its ostensibly stable expression levels. When the distribution of 50-mer overlapping k-mers and their counts are analyzed across 11 isoforms, a complex transcriptional landscape is revealed (as shown in supporting information Fig. S13). For example, isoforms such as GAPDH-201 and GAPDH-211 lack significant unique regions-instead, they share extensive sequences with multiple other isoforms. Although quantification methods like Kallisto, Salmon, Cufflinks, StringTie, and RSEM all suggest that GAPDH-201 is the most abundant isoform, this conclusion is questionable because both GAPDH-201 and GAPDH-211 are similar in length and sequence. This discrepancy likely reflects limitations in the EM or other statistical empirical models that underpin these tools. In contrast, GaussF addresses these challenges by leveraging k-mer counts from minimal and homogeneously shared regions to estimate isoform abundance, potentially overcoming the pitfalls of conventional methods. Consider GAPDH-206, for instance, which features three distinct regions uniquely identifiable along its entire length. These regions capture the complexity introduced by splicing events that extend beyond single exon-intron junctions. To tackle these intricacies, GaussF reprograms the k-mer reference database, establishing a more reliable standard for quantification than methods that consider the full isoform sequence. Additional variability is observed among isoforms that contain single unique regions, such as GAPDH-202, GAPDH-203, and GAPDH-209 (see supporting information, Fig. S14). Notably, no sequencing counts are detected in the unique regions of GAPDH-202 and GAPDH-209. Despite this, Salmon and StringTie are still able to detect GAPDH-202, while Cufflinks and Kallisto report detection of GAPDH-209. Curiously, even though counts are present in the unique region of GAPDH-203, Cufflinks fails to detect it. GaussF addresses these inconsistencies by employing a targeted quantification approach that focuses on isoform-shared specific regions, resulting in more accurate isoform identification and abundance estimation.

### Cross-validation of isoform-level quantification accuracy using the GaussF model

To enhance the validation of GaussF for isoform-level quantification, this study expanded the scope to encompass a larger cohort of human samples, thereby capturing individual variability. It is recognized from prior discussions that isoforms present at low levels are more susceptible to variability than those at high abundance. Consequently, we conducted an extensive profiling of the EMP1-203 isoform across a variety of human colorectal samples (as illustrated in Fig.6a). Intriguingly, within the unique region of this isoform, no instances of 50-mer counts were detected. Upon examination of the adjacent region, which is commonly shared by multiple isoforms, a notable drop in 50-mer counts was observed, suggesting the probable absence of the EMP1-203 isoform in the transcriptomic landscape. The same datasets from 15 individual samples are also analyzed using a variety of quantification techniques i.e., Kallisto, Salmon, Cufflinks, StringTie, and RSEM (as shown in Fig.6b). Surprisingly, the techniques, whether alignment-free (such as Kallisto and Salmon) or alignment-based (like Cufflinks and RSEM), pointed towards a clear detectable abundance that was not trifling. These findings present a notable divergence from the GaussF results, potentially a byproduct of distinct processes used in isoform identification and the assignment of compatible reads to different isoforms. In contrast, the StringTie analysis was congruent with the GaussF results, showing no detectable levels of the EMP1-203 isoform across various human samples. StringTie’s strategy involves the construction of alternative splicing graphs, which is instrumental in its effective reconstruction of isoforms. Similarly, the GaussF approach employs a strategy that echoes this design, especially for the identification of minor isoforms. This parallelism in analytical approach may elucidate the compatibility between StringTie and GaussF findings. Such consistency suggests that these tools may provide a more veracious depiction of isoform presence or absence. In this context, the congruence between StringTie and GaussF lends credibility to their results and underscores the potential for other methodologies to yield false-positive detections due to the intricacies involved in the identification and quantification of complex isoforms. A comparative analysis was performed using various methods to quantify the principal isoform EMP1-201, with every method yielding tangible readouts across all samples (as depicted in Fig.6c). However, the precision of these quantifications is fundamentally intertwined with the successful minimization of multiple biases inherently present in the sequencing data. The GaussF model, which relies on quantification of unique regions and incorporates self-benchmarking, offers a more precise approach to estimate isoform abundance. In addition, an extensive comparison of isoform quantification among various methodologies is examined (as shown in Fig. 6d). This analysis reveals that the quantification abundance of approximately 40% of the isoforms demonstrates notable discrepancies across different methods. Specifically, isoforms that are undetectable according to the GaussF technique can be identified by alternative approaches, whereas certain isoforms clearly identifiable by GaussF are undetectable by these other methods. The Spearman correlation analysis further substantiates the observed inconsistencies across different methodologies when applied to various human samples (as detailed in supporting information Fig. S15). This analysis reveals a method-specific consistency in correlation, wherein a high degree of correlation is maintained across different samples when analyzed using the same technique. This inconsistency poses a significant risk of false positive detections, which in turn, may result in misleading biological interpretations.

**Fig. 6.**
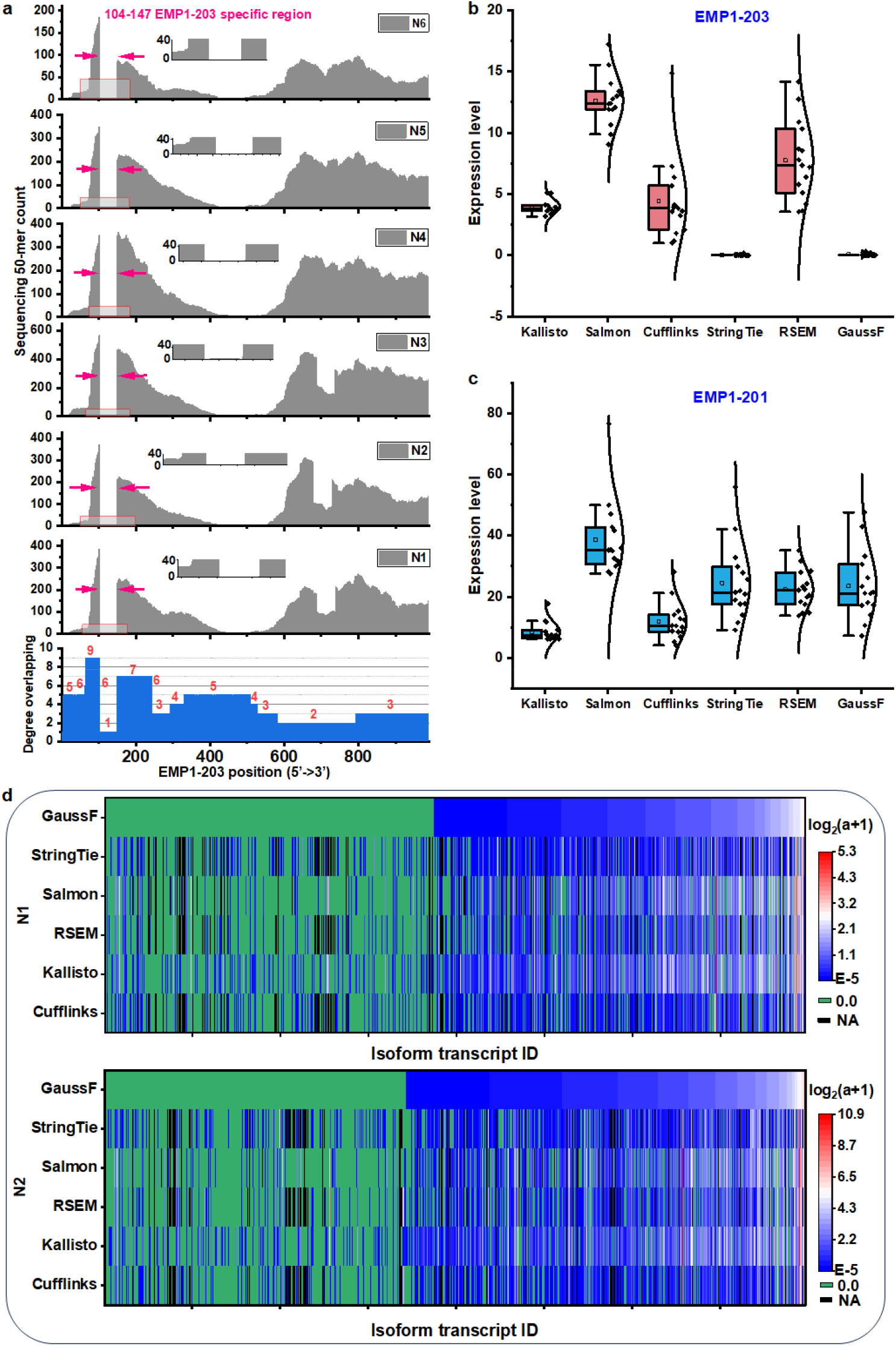
Demonstrating the adaptability of the GaussF algorithm in analyzing various human tissue samples. **a**) The complete 50-mer read count distribution for the EMP1-203 isoform across six human tissue samples is depicted, with annotations highlighting the unique region of this transcript. Each k-mer along EMP1-203 is accompanied by a detailed presentation of its overlap with other isoforms, each region with a precise value. A detailed view of this region is provided in a zoomed-in inset for clarity. **b-c**) The EMP1-203 and EMP1-201 abundance across 15 human samples (including six from (a)) as determined by different methods, including Kallisto (TPM), Salmon (TPM), Cufflinks (FPKM), StringTie (FPKM), RSEM (FPKM), and GaussF (RPKM). **d**) A heatmap comparison presenting the quantification of large-scale isoforms by various methods including Kallisto, Salmon, Cufflinks, StringTie, RSEM, and GaussF across three distinct human tissue samples. The output values are adjusted using a logarithm with base 2 (addition of 1 to the output value before processing). Areas where abundance is null (0.0) or undetectable (na) are specifically highlighted.

Beyond its intrinsic self-benchmarking property, the GaussF algorithm exhibits a sophisticated self-normalizing mechanism that harnesses partially unique regions to depict the overall transcript abundance in RPKM metric. This metric facilitates the adjustment for transcript length, accounting for the contribution of the unique regions to the RPKM value. Such normalization is critical for comparing transcript abundances across different conditions and samples, thus enabling a relatively more accurate and equitable representation of transcript levels. To verify the efficacy of this normalization technique, we identified a circRNA designated as has_circ_0001772, along with several isoforms stemming from the same parent gene, RBM33 (as shown in supporting information Fig. S16a). Notably, the GaussF model introduces a novel capability to align isoforms with their corresponding circular RNAs, thus establishing a direct quantitative link between linear and circular RNA species. This advancement is pivotal in elucidating the complex regulatory landscape encompassing all relevant isoforms (as depicted in supporting information Fig. S16b). Such insights are invaluable for understanding the functional dynamics of RNA molecules and could elucidate their potential involvement in diverse biological pathways. In pursuit of substantiating our observations, the GaussF model was applied to numerous human samples to systematically enumerate all detectable RBM33 isoforms alongside the circRNA (as shown in supporting information Fig. S16c). Despite sample-to-sample variations in isoform abundance, the overarching patterns maintained a remarkable degree of congruence. This underscores GaussF’s capacity to accurately reflect the nuances of alternative splicing phenomena, thereby reinforcing the model’s utility and robustness in transcriptomic analysis.

The GaussF model is acclaimed for its enhanced bias correction measures and serves as an essential benchmark in comparative studies against various transcript quantification models. This study investigates the effectiveness of bias correction in achieving accurate transcript abundance by comparing GaussF’s Gaussian fitting method with other prevalent models based on fragment distribution, GC content, and hexamer usage. Specifically, we focus on the GAPDH-201 isoform as our reference target. In the case of fragment size bias correction, we adjusted the counts of GAPDH-201 fragments to match a global Gaussian distribution, though noteworthy deviations remained (as presented in supporting information Fig. S17a). For GC bias correction, a LOESS smoothing technique was applied to GC-indexed binned data to enhance the fit’s accuracy, providing robust weighting factors across different GC content categories. This method, however, introduced some variability (as shown in supporting information Fig. S17b). The hexamer-based correction addressed bias from preferential hexamer binding by normalizing observed hexamer frequencies to a theoretical uniform distribution, which was used to derive correction factors for fragment counts, detailed in supporting information (Fig. S18c). These diverse bias correction models adopt empirical data to individually calibrate specific bias weighting factors. Contrarily, our self-benchmarking framework leverages theoretical modeling based on Gaussian distributions, utilizing its cumulative distribution function to achieve precise quantification of transcripts such as GAPDH-201 (as illustrated in supporting information Figs. S17d and S17e). This theoretical approach systemically addresses complex biases, providing a more accurate assessment of transcript levels.

To further assess the accuracy of individual bias correction methods, we conducted a comprehensive comparison using four variations of the same Salmon algorithm: uncorrected, only GC bias corrected, only sequence bias corrected, and both sequence-GC bias corrected. Through overlapping plots and correlation analyses (as illustrated in supporting information Fig. S18a), we observed significant discrepancies across the models in isoform-level transcriptome quantification. Despite attempts to align data corrected for GC bias with that corrected for sequence bias, substantial variance persisted, underscoring the challenges in simultaneously addressing all existing biases effectively. Notable examples illustrate these challenges: the specific isoform EMP1-209 (ENST00000542289) was quantifiable only with the GC bias correction model (as shown in Supplementary Fig. S18b), while ACTB-214 (ENST00000645025) abundance dropped to zero after sequence correction. Beyond these extreme outcomes, large variations were common. For instance, the abundance (TPM) of PGK1-203 (ENST00000476531) decreased drastically from 43.7 (uncorrected) to 14.6 after sequence correction, a reduction of nearly two-thirds. Such instances underscore the intricate challenges in achieving accurate transcript abundance estimates, demonstrating the inherent limitations and variable impacts of different bias correction strategies.

## Discussion

The accurate quantification of transcript expression levels, particularly at the isoform level, in sequencing data is confounded by multiple intertwined biases. These biases may stem from a range of factors, including but not limited to GC content, RNA secondary structure, variability in sequencing depth, and the library preparation protocols used. Addressing these biases is crucial for both alignment-based and alignment-free quantification methods to ensure the fidelity of the expression estimates. To mitigate these biases, targeted bias correction methods can be employed, which are designed to neutralize the effects of specific types of distortions within the datasets. The integration of multiple bias mitigation strategies is essential for accurate transcript quantification; however, it presents the challenge of overfitting due to the complex interactions of various sources of bias, which may sometimes be interrelated.^35, 36^ While the conventional quantification methods based on EM flow algorithm have significantly advanced in the modeling and correction of specific biases, they are not comprehensive and simultaneous in addressing observed sources of biases, particularly for those that are more complex and not easily characterized. It is generally accepted that the complete elimination of all biases in transcript quantification is an elusive goal.^37, 38^ Moreover, Traditional methods often presuppose bias weighting factors based on empirical sequencing data, where multiple biases interplay in a complex manner. This greatly complicates efforts to mitigate all biases comprehensively. GaussF distinguishes itself from these standard approaches by simulating an optimal theoretical k-mer distribution model along transcripts to process empirical sequencing data. By applying parametric CDF fitting, GaussF corrects for a whole spectrum of biases. This unique feature positions GaussF as a notable step forward in the realm of bias correction methodologies for sequencing data analysis.

The transcriptome displays non-uniform sequencing coverage, attributed to the overlap of various isoforms and the presence of complex biases within sequencing data. Under conditions of homogenous region, as represented by regions exclusively associated with a single transcript isoform or by regions with a common presence across a consistent set of multiple isoforms, disparities in k-mer distributions can be comprehensively adjusted. Within these segments, erratic coverage is solely attributable to innate sequencing biases, absent the confounding effects of isoform overlap. To effectively mitigate such biases at the isoform level, it is essential to delineate homogeneous regions, with uniform expression patterns, from heterogeneous regions, where expression patterns are influenced by features shared among different isoforms. Consequently, we have selected the as unique as possible region of each isoform for quantification and normalization of data. However, some isoforms lack identifiable unique regions consistently across all isoforms. In the overall alignment with all isoforms, the global transcript for each isoform can be segmented into specific regions based on the components commonly shared among different isoforms. When aiming to compute the sequencing counts according to the global transcript sequence of isoform, we find it necessary to proportionally distribute compatible and ambiguous counts to the isoforms contributing to each locally shared region along the global sequence. Current methods fall short of achieving such level of detail due to the complex interrelationships among isoforms (as shown in Fig.5f), predominantly when dealing with multiple isoforms. An analysis of 20,618 protein-coding genes (from Ensembl BioMart) has yielded a total of 199,384 transcripts at isoform level, averaging approximately 9 isoforms per gene. This finding underscores the intricacy involved in isoform identification and quantification. The EMP1, ACTB, and GAPDH gene serves as the quintessential examples, illustrating the complexity encountered in this process. Indeed, similar challenges are pervasive across nearly all isoforms. To work around this, we have designated the region shared by the least number of isoforms to represent a particular isoform. This implies that the k-mer counts derived from this minimal-isoform shared region could represent the isoform without unique region, but also take the count form other isoforms into account. Moreover, the minimal-isoform shared region strategy offers a significant computational advantage by preemptively excluding regions commonly shared across predominant transcript isoforms. As evidenced by supporting information Fig. S19, adopting this approach substantially reduces the reference scope for 50-mer counting. Such streamlining of the dataset, which potentially contracts the computational domain by an order of magnitude, results in accelerated data processing and more efficient computational resource utilization. This strategy not only increases the tractability of large-scale genomic analyses but also enhances the overall sensitivity and specificity of isoform detection by focusing exclusively on distinct sequence motifs. The targeted approach aligns with the broader objectives of precision genomics, where the expeditious and accurate resolution of complex genetic information is paramount. Moreover, this minimal shared approach is particularly beneficial for downstream biological assays that require highly specific targeting regions, such as RNA interference (RNAi) knockdown assays, antisense oligonucleotide (ASO) knockdown assays, CRISPR interference (CRISPRi) assays, and RNA localization assays. By focusing on the minimal-shared region, GaussF is capable of more accurately predicting gene expression changes and identifying isoform-specific regions for downstream assays.

For such minimal shared regions, the GC content diversity may be insufficient for Gaussian fitting. For example, in ACTB-217 (as illustrated in supporting information Fig. S20a), with only three 50mers, a reliable GaussF fit can’t be achieved. Here, we sum the counts of each kmer instead. Even with enough kmers, GaussF or CDF functions might fail (as seen in supporting information Fig. S20b and S20c), where the GC content distribution shows two peaks or plateaus, preventing a good fit. In these cases, we also use the sum of counts or normalized counts from each 50mer in the minimal shared region to estimate abundance. Our method quantifies 74.4% of isoforms using GaussF fitting (as shown in supporting information Fig. S20d). For the remaining isoforms, where GaussF fitting is unsuitable due to insufficient kmers or abnormal GC distributions, we use the sum of individual kmer counts. A comprehensive analysis of 50mers across the human transcriptome (as illustrated in supporting information Fig. S20e) shows a mean of 371 50-mers per isoform, with a median of 95. The minimal shared region for each isoform typically includes 47 50-mers, the highest percentage (6.3%) among all isoforms. Other significant sets contain 46, 48, 45, and 49 50mers. The largest set includes 43,784 50mers in the NFIA_ENST00000603233.2 isoform, based on data from the Homo_sapiens.GRCh38.cdna.all.fa dataset (www.ensembl.org). The overlapping information across isoforms (as shown in supporting information Fig. S20f) shows a mean overlapping degree of 1.8 and a median of 1. Notably, 70.1% of isoforms have a unique region with an overlapping degree of 1, while 17.5% have an overlapping degree of 2.

The delineation of unique and shared regions is critical for the precise identification of isoforms. To achieve unbiased quantification, normalization of k-mer counts obtained from these regions remains essential. However, the size and position of these regions along the transcript are variable and, in particular, for certain minor isoforms, may be confined solely to the splice junctions between exons.^34^ Estimates of abundance informed by such brief, localized regions are inherently more vulnerable to the non-uniform coverage characteristic of sequencing efforts. To mitigate the impact arising from the brevity of unique regions, one strategy is to augment the size of k-mers such that they consistently span the splice junction sites. This approach serves to enrich the diversification of k-mer sequences and their GC content, as the assortment of sequence variations is directly correlated with k-mer length. However, an excessively long k-mer might overly condense the distribution range of GC content along with its corresponding read counts, which could diminish the fit quality of a Gaussian distribution model. In practice, a 50-mer has been employed for processing various transcript datasets. While this length may not be optimal for every specific transcript, it represents a compromise accepted by most transcripts and accommodates their substantial sequence specificity along with the capacity for normalizing GC content. The GaussF model’s application, incorporating a 50-mer framework for normalization, allows for refined calibration based on these specialized regions, inclusive of splice junctions alone. Nevertheless, such regions defy full-scale extrapolation for normalizing k-mer counts across entire transcripts. In light of these constraints, the refinement of deep learning algorithms tailored for bias alleviation at specific transcript positions is currently underway, signifying an active area of research and development. Although estimates of isoform abundance based on short unique regions may not reflect aggregate transcript values, they constitute a substantial foundation for precise isoform quantification.

Within the domain of transcript abundance estimation, it is pivotal to understand that the modeling principles and methodologies of the GaussF method substantially diverge from those methods, such as Alpine, Salmon or XAEM, particularly concerning bias modeling, correction, the underlying fundamental distributions, and the strategies for estimating abundance.^21, 36, 39–41^ A universally acknowledged strategy in bias modeling involves contrasting empirically observed k-mers or alignments with an unbiased baseline to ascertain instances of over- or under-sampling bias. This strategy, recognized for its efficiency and accuracy, is fundamentally embraced across various methodologies. However, the execution of this strategy diverges significantly between GaussF and other methods. The majority of methods often derive an unbiased background or model directly from the empirical sequencing data itself, inherently assuming that this data, once corrected for perceived biases, can serve as a neutral reference point. Consequently, the reweighted factors applied to account for biases are intrinsically tied to empirical sequencing data. This interdependence introduces a layer of uncertainty to the accurate quantification of isoform abundances, as the process hinges on the assumption that the manipulated dataset sufficiently approximates an unbiased state. In stark contrast, the GaussF method adopts a unique approach by provisioning an ideally unbiased background through the management of GC content distribution. This method assumes that a controlled, ideal data set can provide a more accurate and inherently unbiased baseline for evaluating the presence of sampling bias. We embarked on an in-depth reevaluation of the dataset to meticulously evaluate the obvious presence of 3’ positional bias and to explore the efficacy of the GaussF model in mitigating this particular form of bias. The outcomes of this detailed investigation are comprehensively documented (see supporting information Fig. S21), where a significant reduction in amplitude is calibrated as anticipated. Our rigorous analysis substantiates the manifestation of a pronounced 3’ positional bias, as manifestly indicated by the preferential distribution of 3’ reads (highlighted in Fig. 2b and 2g). Critically, this investigation unveils that the GaussF model possesses the capability to not only identify but also substantially diminish the impact of this predominant bias alongside other potential biases. This revelation underscores the model’s adeptness in counteracting diverse sources of bias, thus bolstering its utility as a potent tool for bias mitigation in sequencing data analysis. Moreover, in the case of isoforms like EMP1-207 and ACTB-217 (see supporting information Fig. S22), which exhibit high abundance within the spectrum of their gene’s isoforms, this notable abundance may originate either from the individual isoforms EMP1- 207 or ACTB-217 themselves or collectively from multiple isoforms of theirs. Our research indicates that when an isoform lacks unique regions, its abundance would tend to be disproportionately high within the total isoform population (as shown Fig. S12 and S13 for GAPDH isoforms, in supporting information). This phenomenon is attributed to the prevalence of compitable and ambiguous reads, which causes these isoforms to be overrepresented when analyzed using traditional sequencing analysis methods, such as Cufflinks, Kallisto, Salmon, RSEM, and StringTie. This methodological approach inadvertently skews the true distribution pattern of isoform abundance, highlighting a pervasive challenge in the quantification of isoforms across the genomic spectrum.

In the GaussF model framework, we introduce an innovative strategy to navigate the transcriptome’s complexity by creating unique k-mer sets for each isoform, delineating minimal-shared regions. These k-mers span from the 5’ to the 3’ end of transcripts, allowing for comprehensive comparative analyses within the transcriptomic spectrum. This involves comparing the sequence of k-mers in an isoform against all transcriptomic k-mers, including those from the same gene. Enhancing our analysis, we define ‘local frequency’ as the occurrence of specific k-mers within an isoform’s unique k-mer set. When k-mer length is too short, local frequency may exceed one, indicating repeated k-mer sequences within the isoform. In contrast, ‘global frequency’ measures the occurrence of an isoform’s k-mers within the entire transcriptome. Remarkably, with 50-mers, local frequency consistently equals one, indicating sequence uniqueness within isoforms. Our method generates a global frequency profile for each transcript, uncovering complex isoform overlaps. By employing 50-mers, we limit global frequency analysis to isoforms from the same gene, identifiable through an isoform’s ID when a k-mer’s global frequency surpasses one. We identify k-mers with the lowest global frequency within an isoform to find minimal shared regions, unique to specific isoforms when global frequency is one. An increase in this minimal frequency signals k-mers shared across isoforms, illustrating transcriptomic intricacy. Minimal shared regions, potentially originating from various isoform combinations, are integrated by the GaussF algorithm despite their discontinuity, offering a comprehensive transcriptome model. This approach streamlines understanding of the transcriptomic complexity and advances genomic research. In certain scenarios, the number of the k-mers within minimal shared regions are remarkably low. A detailed exploration into the transition from a binomial to a normal distribution reveals the complexities associated with these short minimal shared region lengths, often hindering a straightforward mathematical analysis. To address this issue, we introduced a pivotal threshold parameter in the *pipeline_abundance_GaussF_esti_loop.py* script. This parameter specifically targets the k-mer quantity derived from these minimal shared regions. The function of this threshold is to dictate the method used for estimating abundance: when the threshold is set at 10, the script bypasses the GaussF adjustment and directly aggregates the counts to estimate abundance. In contrast, for threshold values greater than 10, the data is adjusted using the GaussF model. This adaptive approach provides a more accurate portrayal of abundance, significantly improving the dependability of the derived conclusions.

## Methods

### Data analysis

1. Data Processing Protocol: A comprehensive software pipeline for the processing of paired-end sequencing data. Initially, FastQC (v.0.11.8) by Babraham Bioinformatics performs a quality assessment of the raw reads. Next, adapter trimming is performed using Cutadapt (v.2.10), followed by further read trimming and cleanup with Trimmomatic (v.0.39). The alignment to the human reference genome GRCh38.p14 (sourced from Ensembl) is conducted via bowtie2 (2.2.5) and STAR (2.7.3a), with subsequent processing by Samtools (v.1.7). The RSeQC package provides quality control post-alignment. Additionally, k-mer segments are aligned to the Ensembl-annotated transcript reference, ensuring accurate transcriptome representation.
2. Cutadapt for adapter removing. During library preparation, adapter sequences-short DNA fragments-are attached to both ends of the DNA strands, which must be removed prior to alignment. For different librarys, the specific adaptators are employed and then removed.
  2.1 N7-N8 library: the following script iterates over samples listed in samplelist.txt, executing Cutadapt with five threads (-j 5) to trim adapter sequences unique to the N7-N8 libraries from both ends of the paired-end reads. The trimmed reads are saved with the suffix _cutadapt_R1.fq and _cutadapt_R2.fq in the specified output directory.

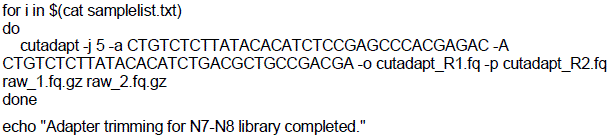

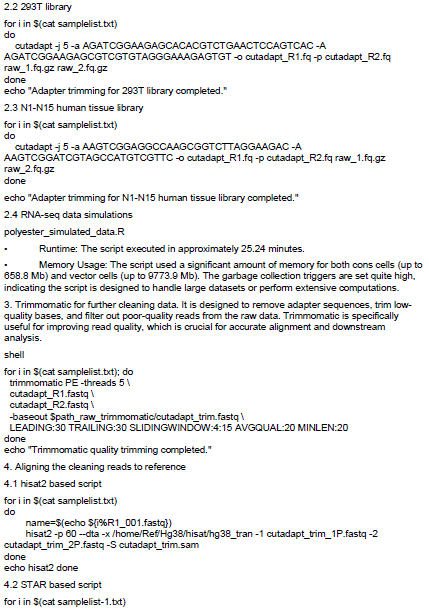

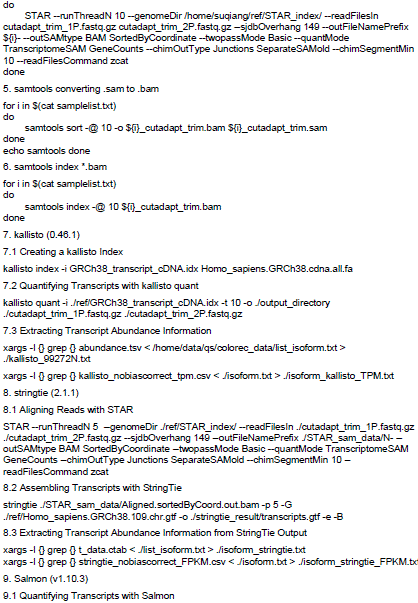

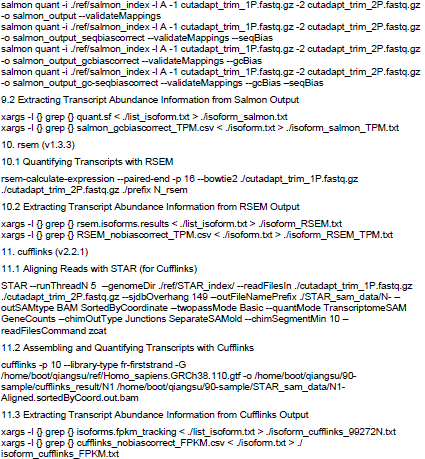

To facilitate the reproduction of analyses from established tools (such as Cufflinks/STAR, StringTie/STAR, Salmon, Kallisto, and RSEM), we have developed a dedicated Snakemake workflow. This workflow is now available in the branch of our GitHub repository: https://github.com/QiangSu/gaussf_pipeline/workflow.

12. GaussF Transcript Quantification Pipeline (https://github.com/QiangSu/gaussf_pipeline)

The GaussF pipeline is designed to accurately quantify transcript abundance at the isoform level using RNA-seq data. The algorithm leverages unique k-mer signatures to overcome common RNA-seq biases, employing a parametric Gaussian model for sophisticated bias correction. This methodology enables precise TPM (Transcript per Million) or RPKM (Reads Per Kilobase of transcript, per Million mapped reads) estimates, facilitating in-depth transcriptomic analyses.

Step 1: Minimal-shared region

The script ***kmer_freq_dist.py*** enumerates specific k-mers for each transcript. The targeted transcript set is defined by the research focus and can be customized based on specific gene lists. Approximately 175,000 transcripts at the isoform level can be processed to assign unique genomic regions. The isoform-specific output will display the following information: Gene Name, which is the name of the gene associated with the k-mers and is typically extracted from the filename of the input CSV file following a predetermined naming convention; Transcript ID, the identifier for the specific transcript from which the k-mers were derived, also extracted from the filename of the input CSV file; Global Frequency, indicating the frequency of the k-mer’s occurrence across all transcripts in the dataset, providing an overall idea of the k-mer sequence’s commonality; Present in Transcripts, an identifier indicating the transcripts that contain the k-mer, presented as either a single transcript ID or a list of IDs, depending on the k-mer’s representation in the data; and Mini Shared Length, which represents the minimum shared length between the input k-mer sequence and any of the transcripts, offering insights into the minimum overlap the k-mer has with known transcripts.

Step 2: k-mer counting

This step involves counting the occurrence of transcript-specific k-mers within quality-controlled fastq datasets, implemented through ***kmer_counter.py***. For each isoform, k-mers and their normalized counts are documented in separate CSV files. These counts are normalized according to base-calling read length, sequencing depth, isoform length, and k-mer length.

Step 3: k-mer merge and normalize

The ***merge_normalize.py*** script merges the normalized k-mer counts from Step 2 with the k-mer reference database. This combines the quantitative count data for each k-mer with its associated reference annotations.

Step 4: GaussF model analysis for unbiased quantification

Finally, the ***gaussf_tpm.py*** script is employed to analyze the normalized k-mer counts derived from the minimal-shared regions. This script utilizes a self-benchmarking algorithm to ensure accuracy. The output of this analysis consists of unbiased TPM and count values for each transcript.

13. A Detailed Procedure for Isoform Alignment and GC Content-Based Modeling

13.1 Isoform Splitting: Divide each isoform sequence into overlapping or non-overlapping 50-nucleotide (50mer) segments. Process all isoforms to create a complete set of 50mers for each.

13.2 Transcriptome-wide 50mer Alignment: Align all generated 50mers against the entire collection of isoform 50mers. This includes matches within the same gene’s isoforms and across different genes’ isoforms.

13.3 Identify Minimal Shared 50mers: Analyze alignment results to find the smallest set of 50mers shared among isoforms of the same gene. This identifies both conserved sequences (present in multiple isoforms) and unique sequences (specific to individual isoforms).

13.4 Characterize Isoform-Specific Regions: Utilize the minimal shared 50mers to define isoform-specific regions and potentially identify functional domains.

13.5 Prepare Initial Data: Create a table for each 50mer in the minimal shared transcript region (identified in Step 3). The table includes the 50mer sequence, a modeling count (initialized to 1), the sequencing count from clean data, the normalized sequencing count (RPKM or TPM), and the GC content.

13.6 Create Example Dataset (Illustrative): Use a simplified dataset (e.g., 10 50mers) to demonstrate the method. This step is primarily for demonstration/testing and might be skipped in a full production pipeline.

13.7 Group by GC Content: Group the 50mers (from the table in Step 5) based on having identical GC content.

13.8 Form Gaussian Distribution: Create a Gaussian distribution based on the GC content groupings (from Step 7), reflecting the expected distribution of GC content.

13.9 Derive Gaussian CDF: Calculate the Cumulative Distribution Function (CDF) from the Gaussian distribution (from Step 8).

13.10 Estimate Parameters: Determine the mean and standard deviation of the GC content distribution from the Gaussian CDF (from Step 9). These parameters are derived from the modeling data’s GC content distribution itself, not from external empirical data.

### Collection of tissue samples

Human colorectal specimens were obtained in compliance with ethical standards from the Sun Yat-sen University Cancer Center and Shenzhen University General Hospital. All participating donors provided informed consent prior to the collection of biopsies, enabling extensive molecular profiling of their transcriptomes for research purposes.

### Cell culture

HEK293T cells were cultivated in DMEM high glucose ((catalog no. SH30022.01, HyClone)), enhanced with 10% fetal bovine serum (catalog no. 10100147, Thermo Fisher), ensuring optimal growth conditions. The culture environment was precisely controlled at 37°C and a 5% CO2 atmosphere with saturated humidity to maintain cell viability.

### Library preparation

The RNA-seq libraries for spike-ins, HEK293T, and colorectal samples were prepared following a standard procedure using the VAHTS Universal V8 RNA-seq Library Prep Kit. This procedure included RNA fragmentation, synthesis of cDNA, end repair, A-tailing, adapter ligation, and library PCR amplification, all culminating in high-throughput sequencing. For spike-in samples, a specialized tagmentation step substituted multiple stages to streamline the library construction process.

### RNA isolation

Total RNA was extracted from cell samples using the RNAiso Plus kit (catalog no. 9109, TaKaRa Biotechnology), adhering strictly to the manufacturer’s protocol. The isolated RNA was dissolved in RNase-free water and quality-checked with a Bioanalyzer RNA picochip. For long-term storage, the RNA was aliquoted and frozen at -80°C to ensure its integrity for future experimentation.

### rRNA depletion

To enhance the resolution of non-ribosomal RNA assessment, we employed the Ribo-off rRNA Depletion Kit (Human/Mouse/Rat) (kit no. N406-01, Vazyme). Through a selective process involving rRNA-targeted probes and enzymatic degradation, this kit efficiently depletes rRNA from the RNA sample. Post-depletion, the purified non-rRNA content underwent quality control checks, readying it for high-fidelity RNA sequencing.

### Spike-in RNA and RNA Circularization

To prepare the spike-in RNA, a single-stranded RNA oligo with a 5’ phosphate and a 3’ OH end was ligated to form a circular RNA. Two spike-in sequences were utilized in the experiment, namely:

Spike-in 1: 5’-phosphate-AAAAAAAAGNTANCTNCGNTTANCACNAGCNCCAAAAAAAA -OH-3’

Spike-in 2: 5’-phosphate-AAAAAAAAGGTAACTGCGNTTANCACNAGCNCCANGAGNAACNACANGAATTCTTTATAAAA AAAA-OH-3’

To create the spike-in RNA, 5 μL of a 10 μM spike-in RNA was added to a 20 μL reaction system containing 1 mM ATP, 2 u/μL RNase inhibitor, 1 μL T4 RNA ligase 1 (ssRNA Ligase) (no. M0204S, New England Biolabs), and 50% PEG8000 with 15% concentration. The reaction was incubated at 25°C for 1-2 hours, followed by termination through boiling for 2 minutes.

### Reverse transcription

The rRNA-free RNA samples underwent reverse transcription using random hexamers and SuperScript™ IV reverse transcriptase (no. 18090200, Invitrogen), following the manufacturer’s instructions.

In a 20 μL reaction system, comprising 100 ng of RNA template, 2.5 μM hexamers, dNTP Mix (10 mM each), and 1 μL (200 units) of SuperScript™ IV, the mixture was incubated at 25°C for 10 minutes, followed by 42°C for 50 minutes. Finally, the reaction was heated to 70°C for 15 minutes to inactivate the reverse transcriptase.

### Tagmentation

For tagmentation, 50 ng of DNA was combined with a 30 μL reaction system consisting of 1× insertion buffer and 2 μL of Tn5-50 adaptor index (10 μM). The mixture was incubated at 55°C for 5 minutes. Subsequently, another 30 μL of 2× Tn5 Digestion Mix (TransNGS® Tn5 DNA Library Prep Kit for Illumina®, no. KP101) was added, and the resulting mixture was incubated again at 55°C for 5 minutes. The resulting tagmented library structure can be represented as follows:

5’-AATGATACGGCGACCACCGAGATCTACAC-i5- TCGTCGGCAGCGTCAGATGTGTATAAGAGACAG-NNNNNN-CTGTCTCTTATACACATCTCCGAGCCCACGAGAC-i7-ATCTCGTATGCCGTCTTCTGCTTG-3’.

### PCR amplication

The library was amplified by PCR using HIFI KAPA master mix (2×) with the following mixture: 25 μL of 2× HIFI KAPA master mix, 10 μL of cDNA, 13 μL of H2O, and 1 μL of either the universal forward or reverse primer (10 μM).

Universal Forward Primer: 5’- AATGATACGGCGACCACCGAGATCTACACCTCTCTATACACTCTT-3’ (:phosphorothioate)

Universal Reverse Primer: 5’-CAAGCAGAAGACGGCATACGAGATGTGACTGGAGTT-3’ (:phosphorothioate)

The PCR reaction was performed in a thermocycler as follows: 95°C for 5 minutes, followed by 10-15 cycles of 95°C for 15 seconds and 60°C for 30 seconds. The amplified library was then purified using 1.8× Ampure XP DNA Beads and resuspended in 20 μL of H2O. One microliter of the library was checked for quality assessment using the Qubit 4 Fluorometer with the dsDNA HS (high sensitivity) Assay Kit from Invitrogen.

### Sequencing

The purified PCR libraries were submitted for sequencing using either the Illumina NovaSeq 6000 (PE150) platform or the MGISEQ-2000 (PE150) platform. The data output was generated according to the specified yield set.

### Statistics

To evaluate the adequacy of linear and Gaussian function fits, we utilized Pearson correlation coefficients and adjusted R-squared values. Additionally, non-parametric tests, specifically the Kolmogorov-Smirnov (K-S) test or Mann-Whitney (M-W) test, were employed to assess the suitability of the models and identify potential deviations between the observed data and theoretical distributions. These tests were conducted with varying sample sizes. The K-S and M-W tests are effective tools for comparing empirical data to theoretical distributions, such as the Gaussian distribution and cumulative distribution. Furthermore, to evaluate associations between variables, appropriate correlation methods were selected based on the data characteristics and the research question. Specifically, for comparing the rank order of transcript abundance estimates, particularly given their often non-Gaussian distribution, Spearman rank correlation was employed to assess monotonic relationships. For assessing linear relationships between other continuous variables where assumptions were reasonably met, Pearson correlation was used.

## Acknowledgments

We acknowledge financial support from the National Key Research and Development Program of China (2023YFA0914904, 2022YFA1105601).This project is in part supported by the Shenzhen-Hong Kong Collaborative Innovation Research (SGDX20201103095404019), National Natural Science Foundation of China (82170070, 82241022), Shenzhen Municipal Research Grant (JCYJ20210324120206017), Yunnan Provincial People’s Hospital Cooperation Project (202201AY070001-224), Guangdong Provincial Key Laboratory of Regional Immunity and Diseases (2019B030301009). We acknowledge financial support from the Long Yi Postdoctoral Special Fund (S249201002). The work presented in the initial submission was performed at Peking University Shenzhen Graduate School.

## Author contributions

Qiang Su and Junmin Quan supervised the initial study. Qizhou Lian supervised all subsequent revisions. Yi Long and Xiaoming Zhou performed structure-related analyses. Qiang Su and Kenneth King-yip Cheng performed data analyses and generated the figures. Yi Long and Qiang Su wrote the manuscript with input from all authors. Deming Gou and Junmin Quan contributed some raw data. All authors read and approved the final version of the manuscript.

## Data availability

The sequencing raw data and processed data have been deposited in the NCBI Sequence Read Archive (https://www.ncbi.nlm.nih.gov/sra) under the accession number PRJNA999048 and PRJNA1051320. The RNA-seq datasets for samples labeled N1 through N15 are available in the Sequence Read Archive under accession numbers SRR25438891, SRR25438890, SRR25438889, SRR25438888, SRR25438887, SRR27182603, SRR27182602, SRR27182605, SRR27182604, SRR27182599, SRR27182598, SRR27182597, SRR27182596, SRR27182595, and SRR27182594, respectively. For the RNA-seq data of N7-N8 with 25-mer and 50-mer spike-ins, replication 1 and replication 2, the SRA accession numbers are SRR25438886 and SRR25438885. Lastly, the SRA accession numbers for the HEK293T total RNA-based RNA-seq, replication 1 and replication 2, are SRR25438884 and SRR25438883. If there are any other relevant data required, please feel free to request them from the authors.

## Code availability

The Python package implementing the GaussF quantification pipeline, gaussf-pipeline, is available under the MIT license on GitHub at https://github.com/QiangSu/gaussf_pipeline. It can be installed directly from PyPI using pip install gaussf-pipeline (see https://pypi.org/project/gaussf-pipeline/). This package provides command-line tools for the complete workflow.

## Supplementary Materials

Figs. S1 to S22

